# CRISPR-based engineering of RNA viruses

**DOI:** 10.1101/2023.05.19.541219

**Authors:** Artem Nemudryi, Anna Nemudraia, Joseph E Nichols, Andrew M Scherffius, Trevor Zahl, Blake Wiedenheft

## Abstract

CRISPR RNA-guided endonucleases have enabled precise editing of DNA. However, options for editing RNA remain limited. Here, we combine sequence-specific RNA cleavage by CRISPR ribonucleases with programmable RNA repair to make precise deletions and insertions in RNA. This work establishes a new recombinant RNA technology with immediate applications for the facile engineering of RNA viruses.

**One-Sentence Summary:** Programmable CRISPR RNA-guided ribonucleases enable recombinant RNA technology.

## Main Text

In 1972, Paul Berg combined restriction endonucleases and DNA ligases to construct the first recombinant DNA molecule (*1*). This work established a new paradigm for DNA manipulation that has altered the way “questions are formulated and the way solutions are sought” (*2*). Initially, DNA manipulations were performed *in vitro*, and the recombinant DNA was delivered to cells for propagation. However, two decades after the first demonstration of recombinant DNA, site-specific nucleases were introduced into cells for *in vivo* editing (*3*). This approach used homing endonucleases to make site-specific DNA breaks that are repaired by cellular DNA repair pathways. But the repertoire of sequences recognized by homing endonucleases is limited, and changing the target specificity is difficult (*4*). Demand for cellular engineering solutions resulted in innovations that made nucleases increasingly more programable (*5, 6*), culminating in the repurposing of CRISPR RNA-guided nucleases for DNA targeting (*7, 8*).

Despite the progress in DNA editing, tools to manipulate RNA without a DNA intermediate remain limited. Although Cas13, a type VI CRISPR RNA-guided endoribonuclease, has been successfully programmed to target cellular transcripts, target recognition also activates a collateral nuclease activity that degrades non-target RNAs (*9, 10*). In prokaryotes, this off-target nuclease activity triggers cell death, providing community-level protection from invading parasites, but the collateral nuclease activity precludes applications that require precision (*11*). Nuclease-inactivated Cas13 (dCas13) has been engineered to stimulate alternative splicing in eukaryotes and deliver RNA-modifying enzymes to specific RNA sequences for nucleobase conversions (adenosine-to-inosine or cytosine-to-uracil) (*12-14*).

Type III CRISPR systems, like type VI (Cas13), target RNA. However, unlike Cas13, the type III CRISPR complexes lack collateral ribonuclease activity (*15, 16*). The absence of collateral cleavage has allowed for precise targeting of cellular transcripts, with minimal off-target effects and cytotoxicity (*17-20*). Here, we combine type III CRISPR-based RNA cleavage with splinted RNA ligation to make programmed deletions and insertions in target RNA sequences. Manipulating RNA is a prerequisite for studying RNA viruses, and we apply CRISPR RNA-guided nucleases to delete or substitute sequences in the genome of the Sindbis virus. This work establishes a new concept for manipulating RNA and demonstrates how this recombinant RNA technology enables rapid and facile engineering of RNA viruses.

### Coupling CRISPR-based RNA cleavage and splint ligation for RNA editing

Type III-A CRISPR complexes comprise five protein subunits (i.e., Csm2-Csm5 and Cas10) that self-assemble in an unequal stoichiometry around a CRISPR RNA (crRNA) (**fig S1A**). The Csm3 protein is an endoribonuclease that forms a helical oligomer along the crRNA and cleaves the bound target RNA in six-nucleotide intervals (**fig. S1B**) (*16*). We hypothesized that the regular cleavage pattern of Csm3 could be leveraged to delete fragments of RNA between each of the cuts (**Fig. 1A**). To test this hypothesis, we cloned, expressed, and purified the type III-A CRISPR complex of *Streptococcus thermophilus* (SthCsm complex) with a crRNA targeting the nucleocapsid (N) gene of severe acute respiratory syndrome coronavirus 2 (SARS-CoV-2) (**fig. S1, C to E**). The purified complex cleaves target RNA with ∼98% efficiency, producing fragments of the expected size (**fig. S1, F and G**).

**Fig. 1.**
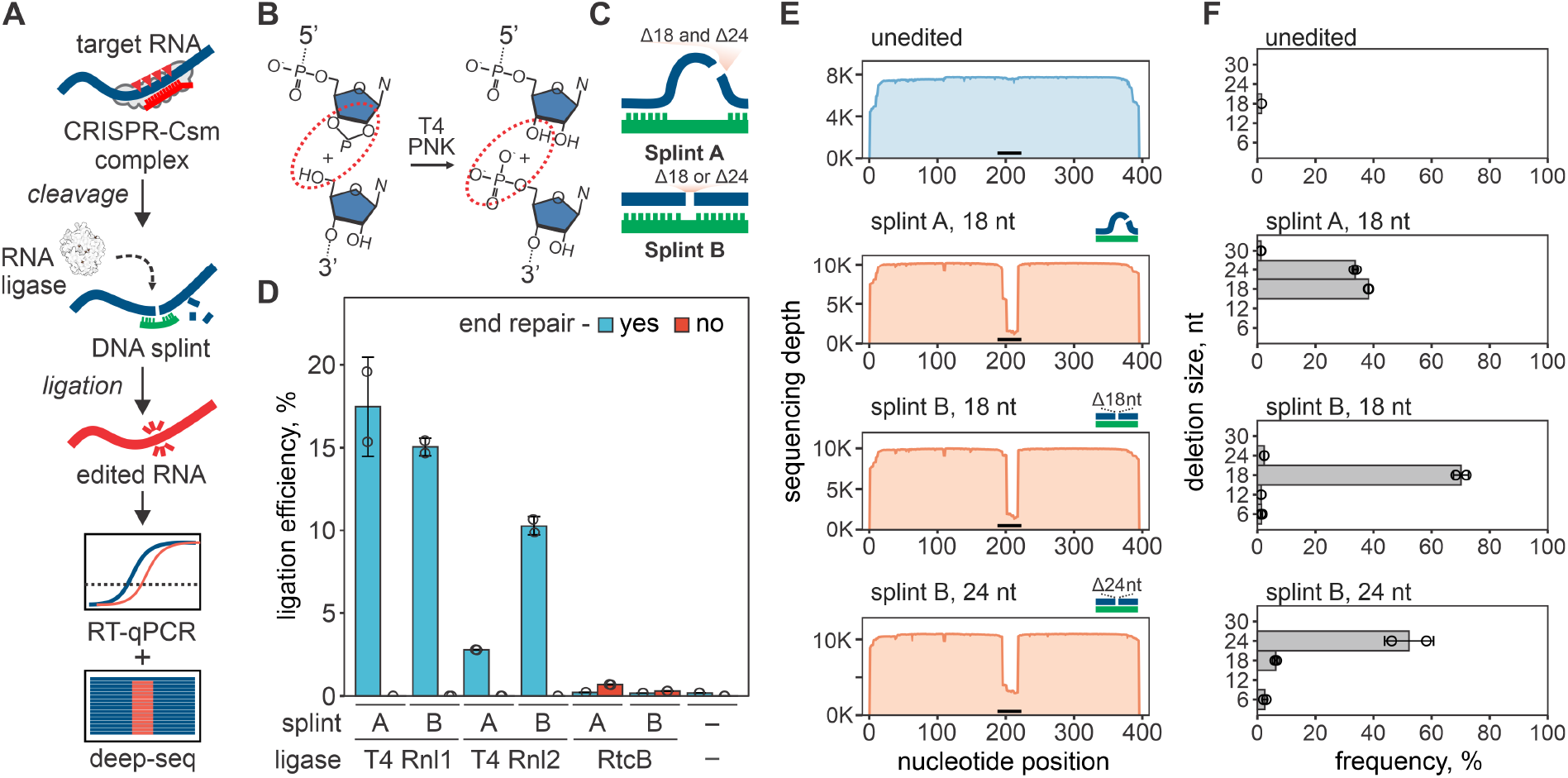
RNA editing with CRISPR-based cleavage and RNA ligation. (**A**) Diagram of the RNA editing with type-III CRISPR nucleases and splinted ligation. Type-III CRISPR complexes cleave RNA in 6 nt intervals (red triangles), cutting out a portion of the target sequence. The resulting fragments are splint ligated to introduce edits. (**B**) SthCsm-mediated cleavage generates 2’3’-cyclic phosphate and 5’-hydroxyl ends (left, substrate for RtcB ligase) that can be converted to 3’-hydroxyl and 5’-phosphate (right, substrate for T4 RNA ligases) using T4 polynucleotide kinase (PNK). (**C**) Two splint designs. Hybridization of the DNA splint A to complementary RNA leaves single-stranded RNA flaps (curved blue lines) that imitate a break in the tRNA anti-codon loop, while splint B mimics nick in double-stranded RNA. (**D**) Comparison of RNA ligation efficiency with T4 RNA ligases (T4 Rnl) and RtcB ligase. Ligation efficiency was measured by performing RT-qPCR across the cut site and quantifying the signal relative to an uncut control (100%). (**E**) Deep sequencing of RNA ligated with splints depicted in (C). Horizontal black bar shows the target site of the SthCsm complex. (**F**) Quantification of editing outcomes in (E).

RNA cleavage by SthCsm generates a 2’,3’-cyclic phosphate and a 5’-hydroxyl at each cut site (*15*). The RtcB ligase of *Escherichia coli* can directly join ends produced by SthCsm cleavage, while RNA ligation by bacteriophage T4 ligases requires a 3’-hydroxyl and 5’-phosphate (*21-23*) (**Fig. 1B**). To facilitate ligation of selected RNA fragments, we used complementary DNA splints that bridge two RNAs. These splints can be designed to imitate different RNA substrates (**Fig. 1C**) (*24*). To identify conditions that result in efficient ligation, we first cleaved *in vitro* transcribed (IVT) RNA derived from the N gene from SARS-CoV-2 using a programmed SthCsm complex. Then we tested three RNA ligases and two DNA splint designs with or without end repair of the cleaved RNA target. As expected, the RtcB directly ligates RNA fragments produced by SthCsm cleavage (0.69 ± 0.01% with splint A), while T4 RNA ligases (T4 Rnl1 and 2) require PNK (polynucleotide kinase) end repair prior to ligation (**Fig. 1D**). RtcB ligation involves fewer RNA manipulations (i.e., no end repair), but ligation with T4 RNA ligases was ∼25-fold more efficient (up to 17.5 ± 3.0% with T4 Rnl1). Therefore, we chose the latter for further experimentation.

To analyze recombinant RNAs in the ligation mixtures, we reverse-transcribed, amplified, and sequenced the targeted regions. Sequencing reads generated with the unedited RNA uniformly cover the length of the amplicon (**Fig. 1E, top**). However, target cleavage followed by splint ligation results in a sharp drop in the sequencing depth at the target site (**Fig. 1E**). Analysis of the sequencing data at the target site identified programmed deletions with high efficiency and accuracy (**Fig. 1F**). Further, we demonstrate that DNA splints can be designed to simultaneously capture multiple editing outcomes (**splint A, Fig. 1F**) or select a specific deletion size (**splint B, Fig. 1F**). This data confirms that target cleavage with SthCsm, followed by splint ligation, results in efficient RNA editing at the specified location.

### Programmed deletions in the RNA virus genome

After validation with *in vitro* transcribed RNA, we next sought to demonstrate how CRISPR-based RNA editing can be applied for the rapid engineering of RNA viruses. As a model, we used Sindbis virus (SINV), an alphavirus with a ∼12.5-kilobase single-strand, positive-sense RNA genome that has a green fluorescent protein (GFP) reporter gene inserted at the 3’-end (**see fig. S2A for details**). We designed a guide RNA to cut out a portion of the *gfp* gene that encodes for tyrosine in position 66 (Y66) of the protein, which would disrupt fluorescence in the encoded protein, such that the edited virus will produce a “dead” GFP (**Fig. 2A**). First, we *in vitro* transcribed the gene for *gfp* and tested the cleavage activity of the *gfp*-targeting SthCsm complex. We sequenced the resulting cleavage products, which identified preferential excision of 6 or 12 nucleotides at the target site (**fig. S3**).

**Fig. 2.**
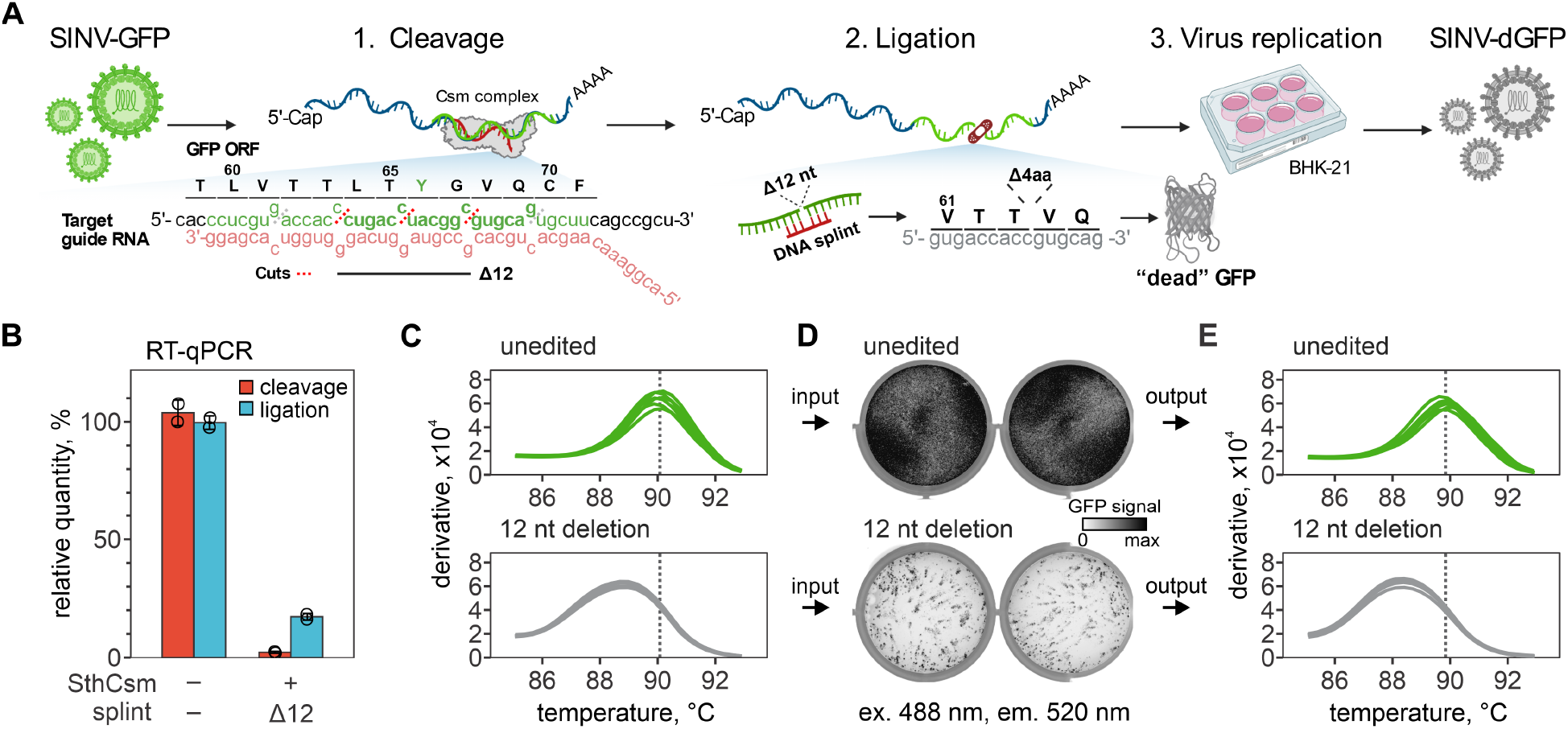
Programmed deletion of specific RNA sequence from the SINV-GFP genome. (**A**) Diagram showing a pipeline for deleting 12 nucleotides from the GFP open reading frame (ORF) of the Sindbis-GFP (SINV-GFP) virus. See Fig. S2A for annotated genome map and Fig. S4A for additional experimental details. The deletion eliminates 4 codons encoding amino acids (L64, T65, Y66, G67) forming the chromophore of the fluorescent protein, which is expected to ablate fluorescence (“dead” GFP, dGFP). (**B**) RNA aliquots collected after the cleavage with the Csm complex (red) and ligation (blue) with T4 RNA ligase were reverse-transcribed and quantified with qPCR. Primers were designed to amplify cDNA across the target site. Relative quantities were calculated by normalizing to the uncut RNA control. (**C**) Melt curve analysis of the qPCR products generated with RNA after ligation in (B). Peaks indicate the melting temperature of the qPCR products. (**D**) BHK-21 cells were transfected with unedited or edited RNA of the SINV-GFP (two replicates each), seeded in 6-well plates, and imaged 24 hours later to capture the GFP signal (excitation 488 nm, emission 520 nm). (**E**) Melt curve analysis of the RT-qPCR products that were generated with RNA extracted from supernatants of BHK-21 cells 24 hours post-transfection. Primers were the same as in (B) and (C).

Next, we moved on to edit infectious viral RNA extracted from the supernatant of infected BHK-21 cells (**fig. S2B**). We incubated viral RNA with a GFP-targeting SthCsm complex and measured the efficiency of cleavage using qPCR (97.8±0.4%, **Fig. 2B**). Cleaved viral RNA was treated with T4 PNK, annealed to a DNA splint designed to select for 12-nucleotide deletion, and ligated with T4 RNA ligase (**Fig. 2B**). Melt curve analysis of the qPCR products spanning the target site identifies a ∼1.5°C shift in the melting temperature of the edited sample, which is consistent with a deletion of the expected size and the faster migration of the PCR product on a polyacrylamide gel (**Fig. 2C, fig. S4, A and B**). Edited or unedited viral RNA was transfected into BHK-21, and cells were imaged the next day. Cells transfected with the edited viral RNA produced significantly less GFP signal compared to the unedited control (*p* = 0.0097; **Fig. 2D and fig. S4, C and D**), while viral RNA concentrations in cell supernatants were nearly identical for edited and unedited viral (*p* = 0.426 for nsP gene and *p* = 0.272 for *gfp* gene, **fig. S4E**). Finally, the PCR performed using a template from the replicating virus suggests programmed deletions at the target site in the *gfp* gene (**Fig. 2E**).

### Sequencing and isolation of edited viruses

To confirm that CRISPR-based RNA editing produces programmed deletions, we sequenced viral genomes extracted from the cell supernatants (**fig. S5**). Sequencing identified a fraction of viral genomes in the unedited control that eliminated the entire *gfp* gene (∼20%, **fig. S5B**). These deletions are also present in the edited sample and accumulate during viral replication in the host because there is no selective pressure on the viral genome to retain the non-essential *gfp* gene. We filtered genomes that retained the *gfp* and quantified deletions of the sequence targeted by the SthCsm complex (**Fig. 3A, fig. S5C**). The 12-nucleotide deletion, which our protocol was designed to select, was the most prevalent editing outcome (34.5 ± 0.9%), but deletions of 18-nucleotides (17.8 ± 2.5%), 24-nucleotides (7.0 ± 2.1%), 30-nucleotides (1.2 ± 0.5%) and 6-nucleotides (2.2 ± 0.8%) could also be identified in the deep sequencing data.

**Fig. 3.**
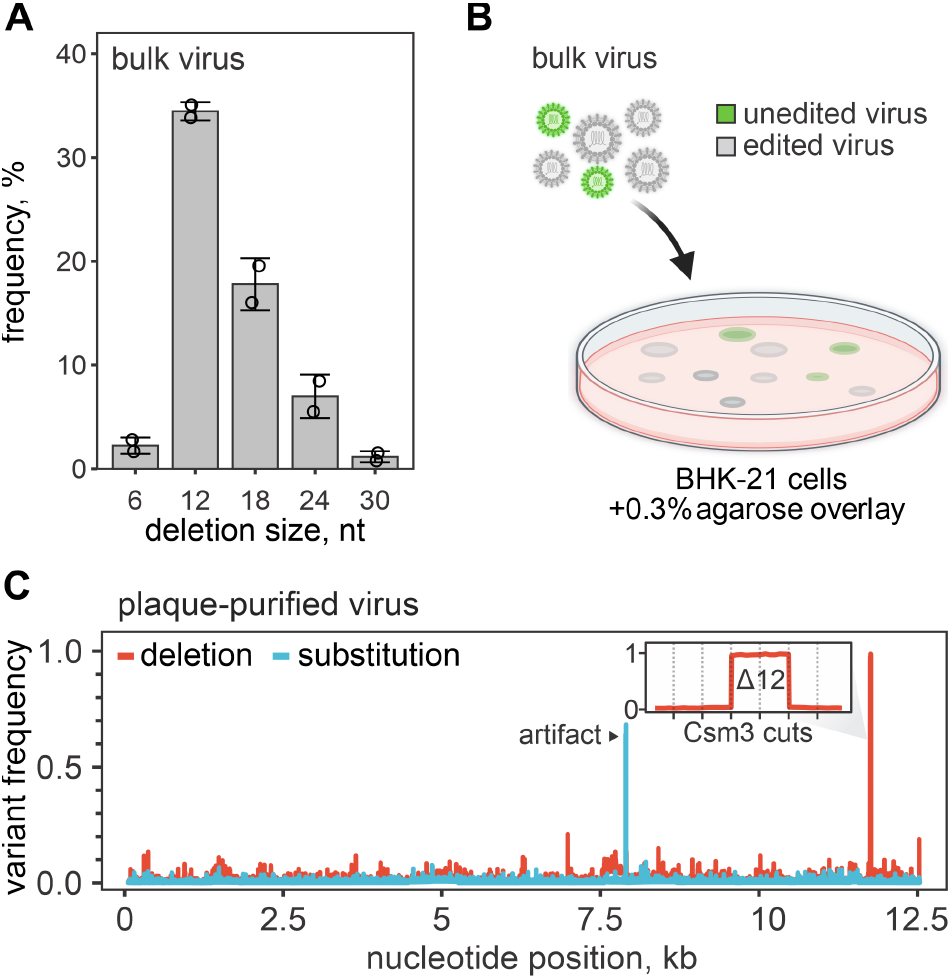
Plaque purification of edited viruses. (**A**) Distribution of deletion sizes in sequencing reads that span the SthCsm target site in bulk edited virus. Data are shown as mean (n = 2) ± standard 15 deviation. Dots show individual replicates. (**B**) Diagram of plaque purification approach used for isolating edited virus clones. (**C**) Frequency of nucleotide variants identified with amplicon sequencing of plaque purified virus. See Fig. S7 for 20 details on sequencing strategy and analysis of other plaque-purified viruses. Black arrow indicates a sequencing artifact associated with amplicon-seq (see fig. S7). The inset is a close-up view of the targeted genomic region. Vertical dotted lines mark 25 predicted cut sites of the SthCsm backbone subunit (Csm3) positioned in 6 nt intervals. The 12 nt deletion was intentionally selected using a DNA splint.

To isolate edited viral clones, we picked plaques from cells transfected with infectious viral RNA and used this as an inoculum to infect individual wells of naïve cells (**Fig. 3B, fig. S6**). We isolated RNA from 11 wells that were positive for cytopathic effect but negative for GFP. The viral RNA was sequenced using amplicon-based genome sequencing (**fig. S7A**). All 11 viral genomes contain programmed deletions at the target site. Four of these genomes contain a 12 nt deletion (**Fig. 3C**), while the remaining seven are a mixture of 12, 18, or 24 nt deletions (**fig. S7**).

### Programmable insertion of new RNA in the viral genome

To demonstrate the versatility of this technology, we designed synthetic RNA and complementary DNA splint to insert a new sequence at the SthCsm cleave site. Instead of SINV-GFP, this time, we used recombinant SINV with a blue fluorescent protein (BFP) gene (SINV-BFP). The nucleotide sequence of SINV-BFP is identical to SINV-GFP except for a single nucleotide substitution (11,762U>C) in the reporter gene that changes tyrosine 66 to histidine (Y66H), resulting in a change in the emission spectrum (i.e., color) of the fluorescent protein. In the synthetic RNA insert, we included a single nucleotide substitution to convert BFP to GFP (11,762C>U) and two additional silent mutations to distinguish the engineered *gfp* from wild-type *gfp* used in other experiments (11,761C>A and 11,764C>U, **Fig. 4A and fig. S8**). We edited the SINV-BFP genome using an SthCsm complex targeting the *bfp* gene, a DNA splint, and a synthetic RNA insert. We transfected the edited RNA and imaged cells the next day. We detected cells that are either blue or green, indicating a mixture of edited and unedited RNA (**Fig. 4, B and C**). To quantify the editing efficiency and to verify that the GFP-positive cells are infected with the edited virus, we sequenced bulk and plaque-purified viruses. These results indicated that 5.1 ± 0.7% of the bulk virus is edited, and the GFP-positive clones contain the intended modification (**fig. S9, 10**).

**Fig. 4.**
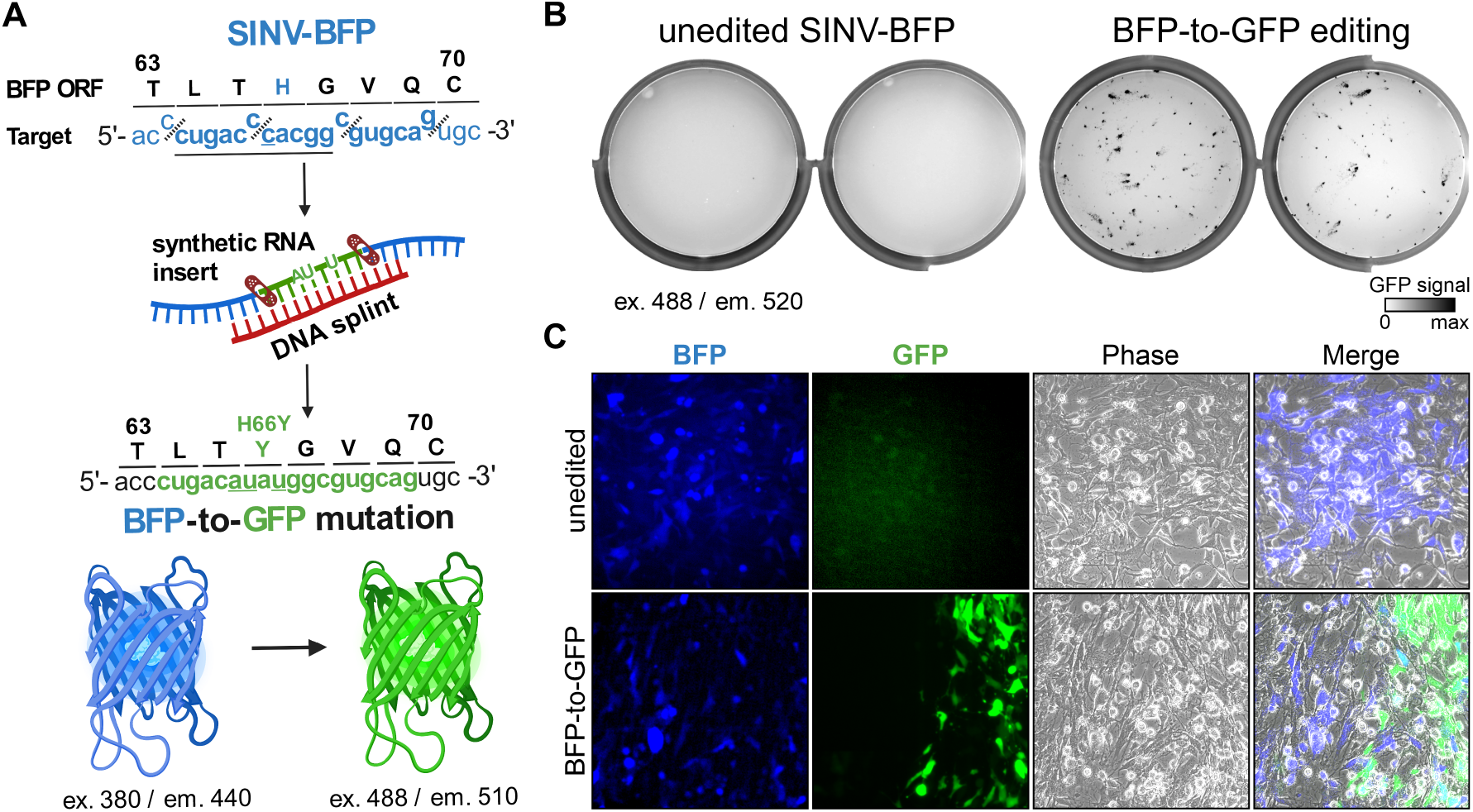
Programmed substitutions in viral genomes. (**A**) Diagram showing editing strategy for substituting 12 nucleotides (underlined) in the blue fluorescent protein (BFP) open reading frame (ORF) of the recombinant Sindbis-BFP (SINV-BFP) virus. The C to U substitution at 11,762 recodes histidine residue for tyrosine (H66Y), which changes the fluorescence from blue to green (BFP-to-GFP). Additional silent substitutions (underlined, 11,761C>A and 11,764 C>U) were included to distinguish edited RNA from the SINV-GFP used for experiments shown in **Fig. 2**. (**B**) BHK-21 cells were transfected with the unedited or edited viral RNA (two replicates each), seeded on 6-well plates, and imaged after 24-hour incubation using 488 nm excitation laser and 520 nm emission filter. This laser and filter setting does not detect BFP (excitation 380 nm, emission 440 nm) in the unedited control, while replication of the edited BFP-to-GFP viruses is visible. (**C**) Transfected BHK-21 cells shown in (B) were imaged with an inverted fluorescent microscope at 20X magnification.

## Discussion

Several years after the Asimolar conference on recombinant DNA in 1975, the first cDNA clones of RNA viruses were created (*25, 26*). Recombinant viruses are a cornerstone of basic research science, leading to fundamental new discoveries in biology, new therapeutics, and vaccines for viral diseases (*27, 28*). Following the outbreak of the severe acute respiratory syndrome coronavirus 2 (SARS-CoV-2), the viral RNA genome (∼30 kb) was reverse-transcribed, and 5 to 12 fragments of the viral genome were cloned into multiple plasmids or an artificial chromosome, which are modified, stitched back together in order to reconstruct the genome, reverse-transcribed and transfected into permissive cells (*29-31*). While recombinant SARS-CoV-2 clones were instrumental in screening antivirals (*32*), studying viral pathogenesis (*31*), and linking genotypes to phenotypes (*33-35*), the reconstitution of mutant viruses is slow and technically challenging, which limits our ability to test new hypothesis (*36*).

Here, we establish a new approach for RNA editing that enables fast and programmable deletions, insertions, and substitutions in any RNA by design (**fig. S11A**). Unlike DNA editing, which often relies on protein-mediated recognition of a specific sequence motif (i.e., PAM), type III CRISPR systems only require complementary base pairing between the RNA guide and the RNA target (*16*), which eliminates additional sequence requirements and improves target site versatility. The protocol described here enables fast and efficient editing of RNA in only three steps (**fig. S11B**). However, the utility of this technology is currently restricted to applications where the editing is performed *in vitro*. Since the majority of plant and animal viruses contain RNA genomes, we anticipate that this is where these tools will have the most immediate significance. Much like DNA editing in cells, edits made in the genomes of RNA viruses are propagated during the infection and inherited by the viral progeny.

We anticipate that *in vivo* RNA editing that relies on the cut-and-repair reaction, similar to those routinely used for DNA engineering, will emerge over time, but applications for base editing might be more immediate. Mutations in type III CRISPR-Cas complexes that eliminate target RNA cleavage but preserve target RNA binding have been well characterized, and the architecture of type III complexes may facilitate the multivalent display of base editors or repair enzymes. Moreover, we expect that diverse type III systems will each have unique properties that are beneficial for applications in editing. Like type II (i.e., Cas9), type V (i.e., Cas12), and type VI (i.e., Cas13), the type III-E systems are composed of a single polypeptide (*37*), and the purified protein can be loaded with synthetic crRNAs *in vitro* (*19, 38*). Loading Cas effectors with synthetic RNA guides was an important advance for DNA editing, and similar advances are anticipated for RNA editing.

## Materials and Methods

### Plasmids

*E. coli* codon-optimized genes for expression of the type III-A Csm complex from *Streptococcus thermophilus* (SthCsm) were synthesized by GenScript and cloned into three different vectors. Genes for SthCsm3 with an in frame N-terminal His6-TwinStrep-SUMO affinity tag, SthCsm4, SthCsm5 were cloned into pRSF-1b (pRSF-1b-SthCsm3,4,5) using NcoI and PacI restriction sites. SthCas10 and SthCsm2 were cloned into pACYCDuet1 (pACYCDuet1-SthCas10-SthCsm2) using NcoI and SacI restriction sites. SthCas6 and CRISPR array containing five repeats and four identical spacers, designed to target the N-gene of SARS-CoV-2, were expressed in the pMA vector (pMA-SthCas6-4xcrRNA-N-gene, GenScript). Cas10 gene with 15HD>HA and 573GGDD>GGAA mutations was synthesized and cloned in pACYCDuet1-SthCas10-SthCsm2 plasmid using NcoI and NotI restriction sites. These mutations inactivate the nuclease and polymerase domains of Cas10, which dramatically improves protein yields (pACYCDuet1-SthCas10^HD/DD^SthCsm2; Fig S3A and B). A CRISPR array containing two repeats and one spacer targeting the *gfp* gene of recombinant Sindbis virus (SINV-GFP) was generated by annealing and extending two partially complementary DNA oligos with Q5 polymerase (NEB) (Table S1). The resulting fragment was cloned using NcoI and MfeI restriction enzymes into pMA-SthCas6-4xcrRNA-N-gene to substitute the CRISPR array. To generate a CRISPR array targeting the *bfp* gene, the plasmid encoding CRISPR targeting *gfp* (pMA-SthCas6-1xcrRNA-GFP) was mutagenized using Q5 site-directed mutagenesis (Table S1). All plasmid sequences were confirmed with whole-plasmid sequencing at Plasmidsaurus (https://www.plasmidsaurus.com/). Plasmid sequences are available at Wiedenheft lab GitHub page (https://github.com/WiedenheftLab/).

### Nucleic acids

DNA oligos were purchased from Eurofins, and RNA oligos from IDT (Tables S1-5). The RNA donor was annealed to a DNA splint at 1:1 ratio in buffer (10 mM Tris-HCl pH 7.8, 50 mM NaCl). Annealing was performed at 95°C for 5 min and ramp down 0.1°C/sec to 25°C (Table S2). *In vitro* transcribed RNA of SARS-CoV-2 N-gene was synthesized with MEGAscript T7 (Thermo Fisher Scientific) using PCR product generated from SARS-CoV-2 WA1 cDNA and primers N0.5_F and N3-5_R (Eurofins) (Table S3). *In vitro* transcribed GFP RNA was synthesized with MEGAscript T7 (Thermo Fisher Scientific) using PCR product generated from pTE3’2J-GFP plasmid and primers T7_SINV_sgmRNA_F and SINV_polyA_R (Eurofins) (Table S3). Transcribed RNAs were purified using the Monarch RNA Cleanup kit (NEB).

### Viruses

Recombinant Sindbis virus expressing GFP (SINV-GFP) was reconstituted from a cDNA copy encoded in pTE3’2J-GFP vector generously provided by Dr. Michelle Flenniken (*39*). RNA (500 ng), *in vitro* transcribed from the pTE3’2J-GFP template, was electroporated into 1 million BHK-21 cells, and the supernatant was harvested 24-48 hours post-electroporation (passage 0). To generate recombinant SINV expressing BFP (SINV-BFP), the *gfp* gene in the DNA template was mutagenized with splice PCR to make 11,762T>C mutation (Table S4). The spliced PCR fragment was cloned in the pTE3’2J-GFP vector using the XbaI restriction enzyme.

### Cell cultures

BHK-21 cells (ATCC, CCL-1) were generously provided by Professor Mark Jutila (Montana State University) and maintained at 37°C and 5% CO2 in Eagle’s Modified Eagle Medium (EMEM, ATCC, cat. #30-2003) supplemented with 10% fetal bovine serum (FBS, ATLAS Biologicals, Lot. #F31E18D1) and 50 I.U./mL penicillin and 50 mg/mL streptomycin. BL21(DE3) protein expression strain was obtained from Thermo Fisher Scientific.

### Protein expression and purification

Expression and purification of the SthCsm complexes were performed according to the following protocol. The expression vectors pRSF-1b-SthCsm3,4,5, pACYCDuet1-SthCas10-SthCsm2, pMA-SthCas6-4xcrRNA-N-gene, and pMA-SthCas6-1xcrRNA-GFP or pMA-SthCas6-1xcrRNA-BFP were transformed in BL21(DE3) cells and grown in LB Broth (Lennox) (Thermo Fisher Scientific) supplemented with 50 μg/mL kanamycin, 34 μg/mL chloramphenicol, and 100 μg/mL ampicillin at 37°C to an OD600 of 0.5. Cultures were then incubated on ice for 1 hour and then induced with 0.5 mM IPTG for overnight expression at 16°C. Cells were lysed with sonication in Lysis buffer (20 mM Tris-HCl pH 8, 500 mM NaCl, 1 mM DTT, 1 mM EDTA), and the lysate was clarified by centrifugation at 10,000xg for 25 mins, 4°C. His6-TwinStrep-tagged protein was bound to a StrepTrap HP column (Cytiva) and washed with Lysis buffer. The protein was eluted with Lysis buffer supplemented with 2.5 mM desthiobiotin and concentrated (10k MWCO Corning Spin-X concentrators) at 4°C. Finally, the protein was purified using a HiLoad Superdex 200 10/300 column (Cytiva) in storage buffer (10 mM Tris-HCl pH 8, 1 mM DTT, 0.1 mM EDTA, 300 mM monopotassium glutamate, 5 % glycerol). Fractions containing the target protein were pooled, concentrated, aliquoted, flash-frozen in liquid nitrogen, and stored at -80°C.

### SINV genomic RNA (gRNA) purification

Six T300 flasks of confluent BHK-21 cells were infected with SINV-GFP or SINV-BFP viruses (passage 0). Viral stock (50 μL) was diluted in 50 mL of PBS supplemented with 3% FBS, 0.5 mM MgCl_2_, and 0.9 mM CaCl_2_. Media was discarded, and 8 mL of diluted viral stock was added to each flask. Cells were incubated for 1h (37°C, 5% CO2), PBS was removed, and 40 mL of fresh EMEM supplemented with 5% FBS, 50 I.U./mL penicillin and 50 mg/mL streptomycin was added. Supernatants were collected 24 hours post infection (hpi), combined and spun down for 10 min at 3,000 g to remove cell debris. The supernatant was filtered through a 0.45 μm filter and ultracentrifuged in SW32 rotor at 100,000xg for 1 hour at 4°C. The supernatant was discarded, and the pellet was dissolved in 300 μL of PBS. Ten volumes of TRIzol Reagent (Invitrogen) were added to one volume of viral suspension in PBS, mixed, and incubated for 10 min at room temperature. Then, 0.2 volume of chloroform per 1 volume of TRIzol was added, mixed, and centrifuged at 5,000xg, for 10 min at 4°C to separate aqueous phase. The aqueous phase containing RNA was transferred into a new tube, mixed with one volume of 98% ethanol, and purified with Monarch RNA Cleanup kit (NEB). RNA concentration was measured using Nanodrop. This protocol yields ∼80-90 μg of RNA, which is 0.3-0.35 μg per mL of supernatant. The extracted viral RNA is infectious, with 8.3 ± 0.6 × 10^4^ plaque-forming units per microgram of the stock (**fig. S2B**).

### Editing of in vitro transcribed RNA

For the experiment shown in Fig 1C, IVT RNA of SARS-CoV-2 N-gene (0.5 μM) was mixed with SthCsm (1 μM) in a reaction buffer (20 mM HEPES pH 8.0, 50 mM KCl, 1 mM Mg(CH3COO)_2_, 0.1 mg/mL BSA) and incubated for 1 hour at 37°C. The RNA cleavage products were purified using Monarch RNA Cleanup kit (NEB) and divided into two parts. One part of RNA was treated with T4 PNK (NEB) in a reaction buffer (50 mM Tris-HCl pH 7.5, 10 mM MgCl2, 1 mM ATP, 10 mM DTT) for 1 hour at 37°C. The remaining RNA was incubated in the same buffer but without the enzyme. Next, RNA was purified with Monarch RNA Cleanup kit (NEB). PNK-treated and non-treated RNAs were annealed to splints (1:1 ratio) in a reaction buffer (50 mM Tris-HCl pH 7.5, 10 mM MgCl2, 1 mM DTT) for 5 min at 95°C, followed by a ramp down of 0.1°C/sec to 25°C. Then, three different ligases were tested to repair RNA. T4 RNA ligase 1 was mixed with splinted RNA in a 1x T4 Rnl buffer (NEB) supplemented with ATP (1 mM) and PEG 8000 (10%) and run for 1 hour at 25°C. The reaction with T4 RNA ligase 2 was performed in 1x T4 Rnl2 (NEB) supplemented with PEG 8000 (10%) and run for 1 hour at 25°C. The reaction with RtcB was performed in 1X RtcB reaction buffer (NEB) supplemented with GTP (0.1 mM) and MnCl2 (1 mM), and PEG8000 (10%) and run for 1 hour at 37°C. The reactions were diluted 10 times and used for RT-qPCR.

The experiment shown in Fig 1, D-F, was performed as described above with slight modifications. All RNA was T4 PNK-treated after the cleavage with SthCsm. T4 RNA ligase 1 was used to ligate RNA annealed to splint A. To ligate RNA hybridized with splint B, T4 RNA ligase 2 was used. Ten-fold diluted reactions were used for RT-qPCR and amplicon sequencing with Oxford Nanopore.

### Editing of viral RNA

For viral RNA editing (Figs. 2-4), 1.64 μg of viral gRNA (∼0.04 pmol based on MW of the full genome) was mixed with SthCsm-Cas10^HD/DD^ (4 pmol) in a 10 μL reaction in buffer (50 mM Tris-HCl pH 7.5, 1 mM DTT, 1 U/μL RNase Inhibitor (NEB)) and incubated for 1 hour at 37°C. The RNA cleavage products were purified using Monarch RNA Cleanup kit (NEB) and used for a one-pot reaction of end-prep and splint annealing. RNA was mixed with 5 U of T4 PNK (NEB), DNA splint (0.4 pmol) in 10 μL reaction in buffer (50 mM Tris-HCl pH7.5, 1 mM DTT, 1 mM MgCl2) supplemented with 1 mM ATP and 1 U/μL RNase Inhibitor (NEB). Reaction was incubated for 1 hour at 37°C. After the incubation, 5 μL of a mix containing 5 U of T4 RNA Ligase 2 (NEB), 1 mM ATP, 3 μL of 50% PEG 8000, 1 U/μL RNase Inhibitor (NEB) in reaction buffer (50 mM Tris-HCl pH7.5, 1 mM DTT, 1 mM MgCl2) was added directly to the previous reaction. Ligation reactions were incubated 1 hour at room 25°C. Ligation reaction (10 μL) was mixed with 10^6^ BHK-21 cells in 100 μL of Ingenio® Electroporation Solution (Mirus) and transferred to a 0.2 cm cuvette (Mirus). The electroporation was performed using Ingenio® EZporator® Electroporation System (Mirus) by pulsing with 150V and 950-1050 μF. Immediately after electroporation, 500 μL of EMEM (ATCC) media supplemented with 10% FBS (no antibiotics) was added directly in the cuvette. Cell suspensions were transferred to tissue culture plates with addition of more media and incubated with 5% CO_2_ at 37°C.

### Fluorescence imaging

Viral infections were imaged with Amersham Typhoon 5 scanner (Cytiva) with default Cy2 (excitation 488 nm, emission 520 nm) setting. Images of cells infected with SINV BFP-to-GFP edited virus were captured with Nikon Ti-Eclipse inverted microscope (Nikon Instruments) equipped with a SpectraX LED excitation module (Lumencor) and emission filter wheels (Prior Scientific). Fluorescence imaging used excitation/emission filters and dichroic mirrors for GFP and DAPI (Chroma Technology Corp.). Images were acquired with Plan Fluor 20Ph objective and an iXon 896 EM-CCD camera (Andor Technology Ltd.) in NIS-Elements software. Nucleic acid gels were stained with SYBR Gold (ThermoFisher Scientific) and imaged using Cy2 setting with Amersham Typhoon 5 scanner (Cytiva). Protein gels were stained with Coomassie stain (in-house), washed and imaged with IR-short default setting with Amersham Typhoon 5 scanner (Cytiva).

### Plaque purification

The cells electroporated with edited viral RNA were serially diluted, and 10,000 cells were seeded on a monolayer of BHK-21 cells in a 100 mm Petri dish. The cells were incubated (37°C, 5% CO_2_) for one hour. Next, the media was removed, and media supplemented with 0.3% agarose was overlayed. Petri dishes were imaged 16-18 hours post-infection using Amersham Typhoon 5 scanner (Cytiva). The individual plaques were picked using sterile tips under a light microscope and inoculated in naive BHK-21 cells for propagation.

### RT-qPCR

RNA was extracted from 140 μL of cell supernatants with QIAamp Viral RNA Mini Kit (QIAGEN, # 52906) and eluted in 60 μL of buffer AVE (QIAGEN). RNA ligations were diluted tenfold and used for reverse transcription without extraction. The reverse-transcription (RT) reactions were performed with 2X LunaScript® RT SuperMix Kit (NEB) with 5 μL of RNA input in 10 μL total volume. The RT reactions containing cDNA were diluted 100-fold and 5 μL was used for qPCR reactions with 2X Universal SYBR Green Fast qPCR Mix (ABClonal). Each qPCR reaction contained 5 uL of cDNA, 4.2 μL of Nuclease-free Water, 0.4 μL of 10 mM Primers (Table S4), and 10 μL of 2X mastermix. Nuclease-free water was used as negative template control (NTC). Three technical replicates were performed for each sample.

Amplification was performed in QuantStudio 3 Real-Time PCR System instrument (Applied Biosystems) as follows: 95°C for 3 min, 40 cycles of 95°C for 5 s and 60°C for 30 s, followed by Melt Curve analysis. Results were analyzed in Design and Analysis app at Thermo Fisher Connect Platform.

### Nanopore sequencing

Edited IVT RNA of SARS-CoV-2 N gene was sequenced using Oxford Nanopore with Ligation Sequencing Kit (SQK-LSK109). RNA was reverse transcribed using LunaScript® RT SuperMix Kit (NEB), and cDNA was used for PCR with primers nCoV-96_L/nCoV-96_R (Table S4) and Q5 polymerase (NEB). DNA was purified using magnetic beads (Omega Bio-tek, M1378-01) and used to prepare sequencing libraries as described in SQK-LSK109 protocol using Native Barcoding Expansion 1-12 (EXP-NBD104). Resulting library (∼20 ng) was sequenced on the Nanopore MinION with R9.4.1 flow cell. The sequencing run was performed in the high-accuracy base calling mode in the MinKNOW software.

Products of RNA cleavage with *gfp*-targeting SthCsm were sequenced with Oxford Nanopore using direct cDNA sequencing kit (SQK-DCS109). After incubation with SthCsm complex, RNA cleavage fragments were purified with Monarch RNA Cleanup kit (NEB). Then, RNA was treated with T4 PNK (NEB) for 1 hour at 37°C and purified with RNAClean XP beads (Beckman Coulter). The RNA ends were extended using *E. coli* polyA polymerase (NEB), and RNA was cleaned up again using RNAClean XP beads (Beckman Coulter). ∼100 ng of poly(A) RNA was used as an input to prepare cDNA for nanopore sequencing as instructed in SQK-DCS109 protocol using Native Barcoding Expansion 1-12 (EXP-NBD104). ∼200 fM of library was loaded on the Nanopore MinION (R9.4.1 flow cell). The flow cell was primed, and library was loaded according to the Oxford Nanopore protocol (SQK-LSK109 kit). The sequencing run was performed in the high-accuracy base calling mode in the MinKNOW software.

The bulk edited SINV genomes were sequenced with Oxford Nanopore using a direct cDNA sequencing kit (SQK-DCS109). Plaque-purified clones were sequenced with amplicon sequencing using Ligation Sequencing Kit (SQK-LSK114). To perform amplicon sequencing, RNA was reverse-transcribed using LunaScript® RT SuperMix Kit (NEB). Diluted cDNA (100-fold) was used for two multiplex PCR reactions with Q5 polymerase (NEB) with two primer pools (Table S5). The primers were designed using the Primal Scheme web tool (http://primalscheme.com), synthesized (Eurofins), and diluted as previously described (*40*). After amplification, two multiplex PCR reactions for each sample were combined and purified using magnetic beads (Omega Bio-tek, M1378-01). ∼130 ng of DNA was used to prepare sequencing libraries as described in SQK-LSK114 protocol using Native Barcoding Kit 24 V14 (SQK-NBD114.24). ∼20 fM of the library was loaded on the Nanopore MinION (R10.4.1 flow cell, 260 bps). For direct cDNA sequencing, 420 μL of viral supernatant was extracted with QIAamp Viral RNA Mini Kit (QIAGEN, # 52906) and eluted in 30 μL of buffer AVE (QIAGEN). Extracted RNA (7.5 μL) was reverse-transcribed, and the second strand was synthesized as instructed by the manufacturer’s protocol (SQK-109). The resulting double-stranded DNA was used for sequencing library preparation with Native Barcoding Kit 24 V14 (SQK-NBD114.24). The flow cell was primed, and sequencing libraries were loaded according to the Oxford Nanopore protocol (SQK-LSK114). The sequencing run was performed in the fast base calling mode in the MinKNOW software. Sequencing data (.fast5 files) was basecalled with the super-accuracy model (dna_r10.4.1_e8.2_260bps_sup.cfg) using guppy_basecaller and demultiplexed with guppy_barcoder (both from Oxford Nanopore). Basecalling was performed on the Tempest High Performance Computing System, operated and supported by University Information Technology Research Cyberinfrastructure at Montana State University.

### Sequencing data analysis

Nanopore sequencing reads generated with R9.4.1 flow cells were basecalled in high accuracy mode in MinKNOW, and sequencing errors were corrected using the isONcorrect package (*41*). Nanopore sequencing reads generated with R10.4.1 flow cells were collected at 260 bps speed and basecalled in super accuracy mode with guppy_basecaller using dna_r10.4.1_e8.2_260bps_sup.cfg configuration. Sequencing reads were aligned to the reference using *minimap2* (v2.17 -r954-dirty) with Nanopore preset (-ax map-ont setting). Alignments were converted to BAM format, sorted, and indexed using *samtools* v1.13. Sequencing depth was computed with the *samtools depth* program. Sequencing reads generated with amplicon-seq were trimmed to remove primer binding regions using the *samtools ampliconclip*. Genome-wide deletion and substitution frequencies were quantified and normalized by calculating frequency of each nucleotide variant at every position in the viral genome using *pileup* function in *Rsamtools* v2.8.0 package in R. Deletion sizes were identified using the extract-junctions.py script (github.com/hyeshik/sars-cov-2-transcriptome/blob/master/nanopore/_scripts/extract-junctions.py) (*42*). Deletions were quantified and filtered using custom R scripts. Reads with complete deletions of *gfp* or *bfp* were identified, and resulting read id lists were used to filter alignment files (.BAM) using the *samtools view*. Deletion counts at the SthCsm cleavage site were normalized to the number of reads that span the target region, which was computed by extracting read information from BAM files using the *bamtobed* function in *bedtools* v2.30.0 package and filtering reads by start and end coordinate. All plots were created using *ggplot2* v3.3.5 package in RStudio software v2023.03.0.

### Statistical analysis

RNA editing experiments were performed in two replicates. All RT-qPCR reactions were performed in two technical replicates. Statistical comparisons in **Fig. S4D** and **E** were performed in R with Welch’s unequal variances *t*-test using *t*.*test* function from *stats* package. Significance levels: \**p* < 0.05, \*\**p* < 0.01, \*\*\**p* < 0.001, *ns* – non-significant.

## Acknowledgments

We thank Dr. Michelle Flenniken for providing the cDNA clone of the Sindbis-GFP virus (pTE3’2J-GFP) and the related protocols. We thank Dr. Mark Jutila for kindly providing BHK-21 cells. We thank Dr. Matthew Taylor for the generous use of the fluorescent microscope. Some diagrams and schematics were created with BioRender.com. Base-calling of Nanopore sequencing data was performed with the Tempest High Performance Computing System, which is maintained by University Information Technology Research Cyberinfrastructure at Montana State University.

## Author contributions

Conceptualization: B.W., A. Nemudryi, A. Nemudraia, and J.E.N. Methodology: A. Nemudryi and A. Nemudraia. Investigation: A. Nemudryi, A. Nemudraia, J.E.N., A.M.S., and T.Z. Formal analysis: A. Nemudryi and A. Nemudraia. Visualization: A. Nemudryi and A. Nemudraia. Supervision: B.W., A. Nemudryi, and A. Nemudraia. Funding acquisition: B.W. and A. Nemudryi. Writing - Original Draft: B.W., A. Nemudryi, and A. Nemudraia. Writing - Review & Editing: B.W., A. Nemudryi, and A. Nemudraia.

## Data and materials availability

All sequencing data and code for data analysis are deposited to NCBI Sequence Read Archive (SRA) under a single Bioproject and the Wiedenheft lab GitHub page (https://github.com/WiedenheftLab/). Data will be released upon peer-review and publication. Purified proteins, plasmids, and viruses are available from the corresponding author upon request.

