## Supplemental Figures and Tables for "CRISPR-based engineering of RNA viruses"

**The PDF file includes:**

Figs. S1 to S11
Tables S1 to S5

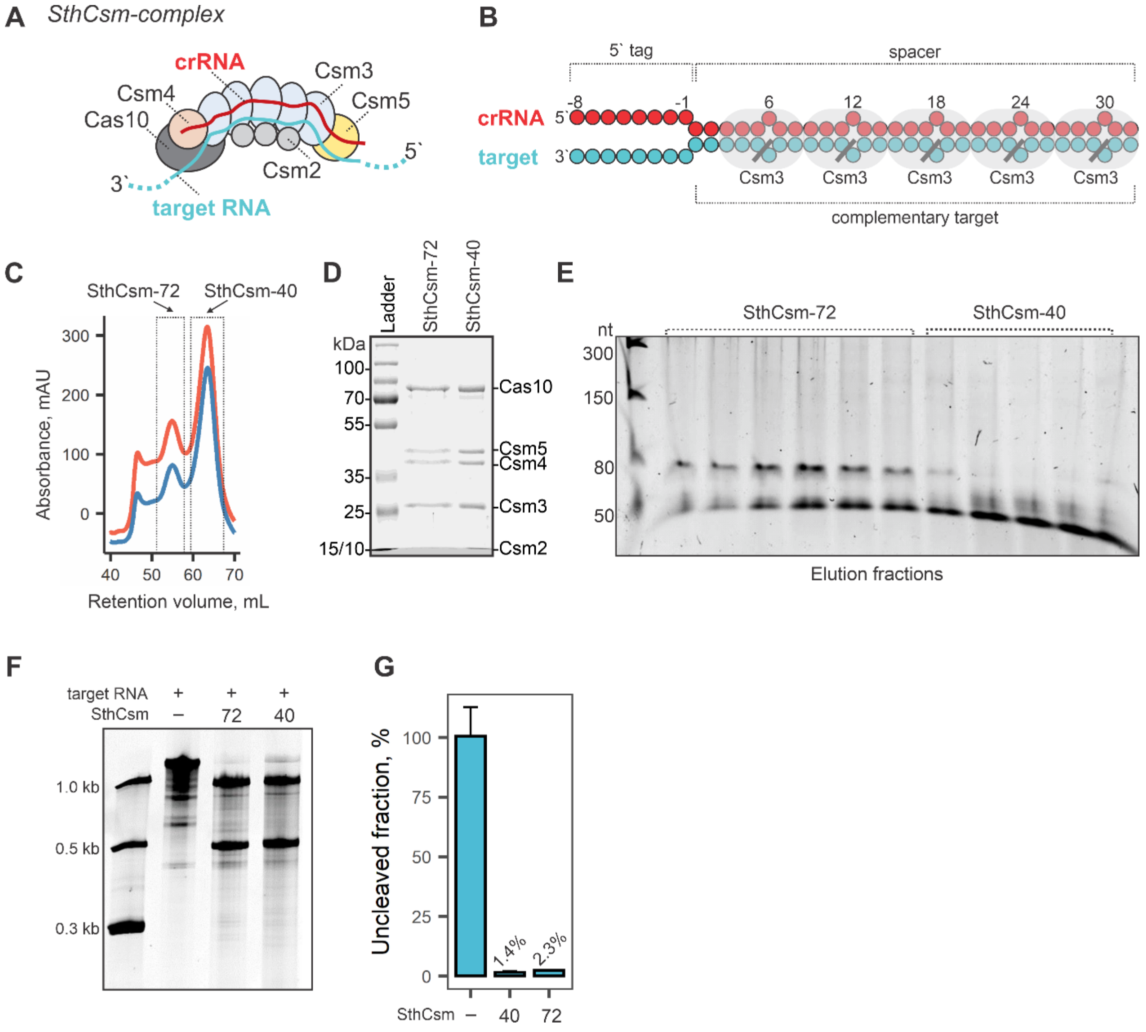


**Fig. S1. Purification of type III CRISPR complex of *Streptococcus thermophilus* (SthCsm).**

(**A**) Diagram of tertiary SthCsm complex with crRNA and target RNA. The complex comprises five protein subunits (i.e., Csm2-5 and Cas10), which self-assemble along the crRNA (red). Csm3 subunits in the backbone of the complex cleave target RNA (blue)

(**B**) Diagram of the crRNA (red) bound to the target RNA (blue). Circles depict ribonucleotides. Multiple copies of the Csm3 subunit in the complex backbone cut bound RNA at six-nucleotide intervals (diagonal black lines).

(**C**) Size-exclusion chromatography (SEC) of affinity-purified SthCsm complex. SEC was performed using a S200 10/300 size-exclusion column (Cytiva). SthCsm complex eluted in two peaks that correspond to different crRNA lengths, i.e. SthCsm-72 (fractions #6-15) and SthCsm-42 (fractions #20-32) (*16*).

(**D**) SDS-PAGE of purified SthCsm-complexes (72 and 42) (~3 ug/lane).

(**E**) RNAs were extracted from SEC fractions containing SthCsm complex shown in (C) and loaded on 12% UREA-PAGE.

(**F**) Cleavage of IVT target RNA with SthCsm-complexes. Reactions were incubated for 1 h at 37°C, purified with Monarch RNA Cleanup kit (NEB) and loaded on 10% UREA-PAGE.

(**G**) Cleavage efficiency was quantified with RT-qPCR across the cut site. Data is shown as the mean of two replicates ± one standard deviation.

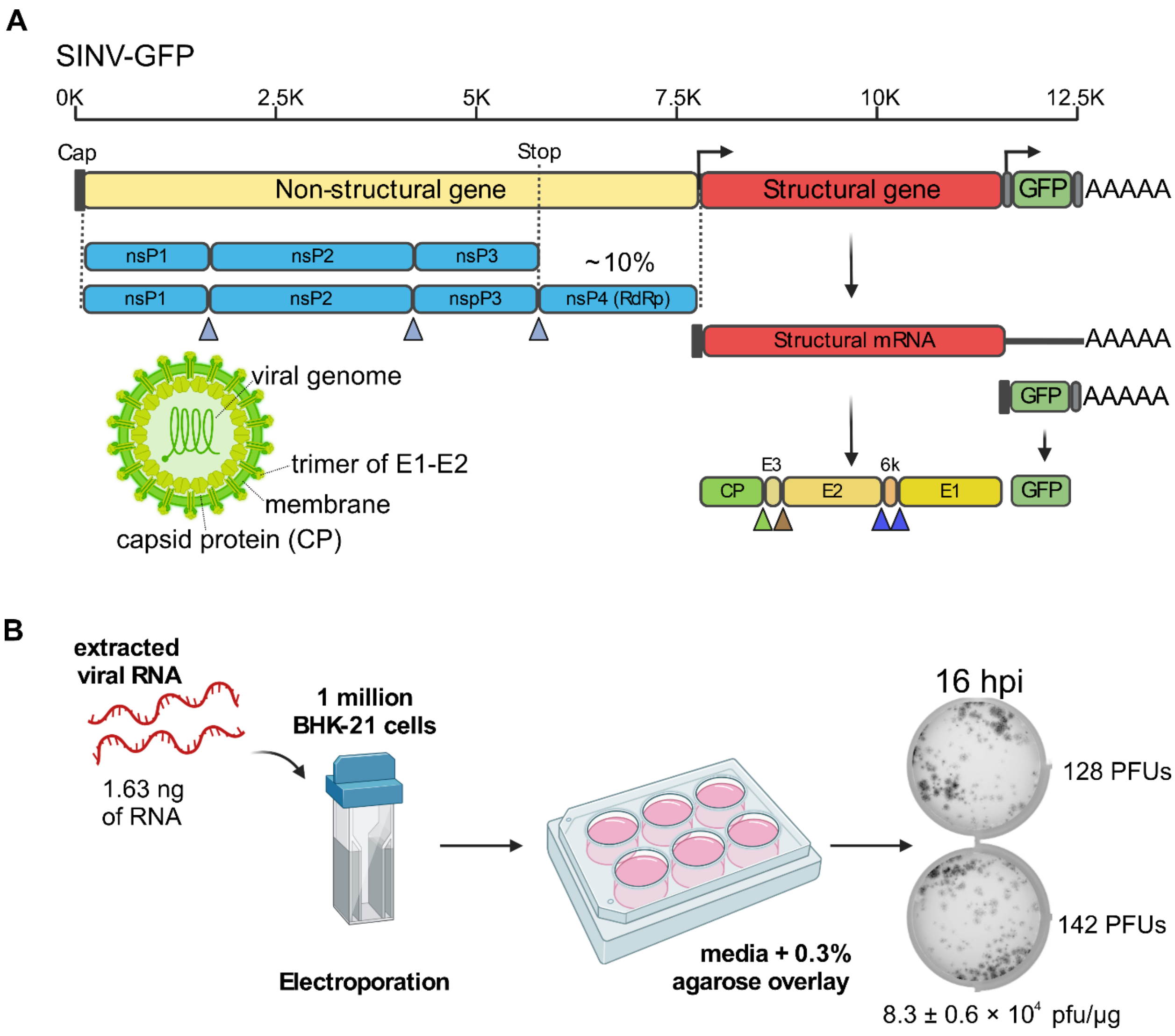


**Fig. S2. Architecture of recombinant Sindbis GFP virus (SINV-GFP) and tittering viral RNA.**

(**A**) Sindbis virus (alphavirus genus, *Togaviridae* family) has a ~12.5-kilobase single-strand, capped and polyadenylated positive-sense RNA genome with two open reading frames that encode non-structural (nsP1-nsP4) and structural proteins (CP, E1-E3, 6K). Triangles indicate proteolysis sites. The structural gene is expressed from an internal promoter, which generates a subgenomic messenger RNA (sgmRNA). The recombinant Sindbis-GFP virus (SINV-GFP) used in this work has a *gfp* gene integrated at the 3'-end of the genome, which is expressed from an internal promoter and makes additional sgmRNA.

(**B**) Viral RNA, concentrated and extracted from supernatants of infected cells, was diluted and transfected in BHK-21 cells (see *Methods*). Cells were seeded in a 6-well plate, and after 1 h incubation at 37°C with 5% CO2 media was replaced with fresh media supplemented with 0.3% agarose. After 16 h incubation, cells were imaged with Amersham Typhoon 5 scanner, and viral plaque forming units (PFUs) were quantified. Resulting numbers were used to calculate titer in the undiluted RNA stock.

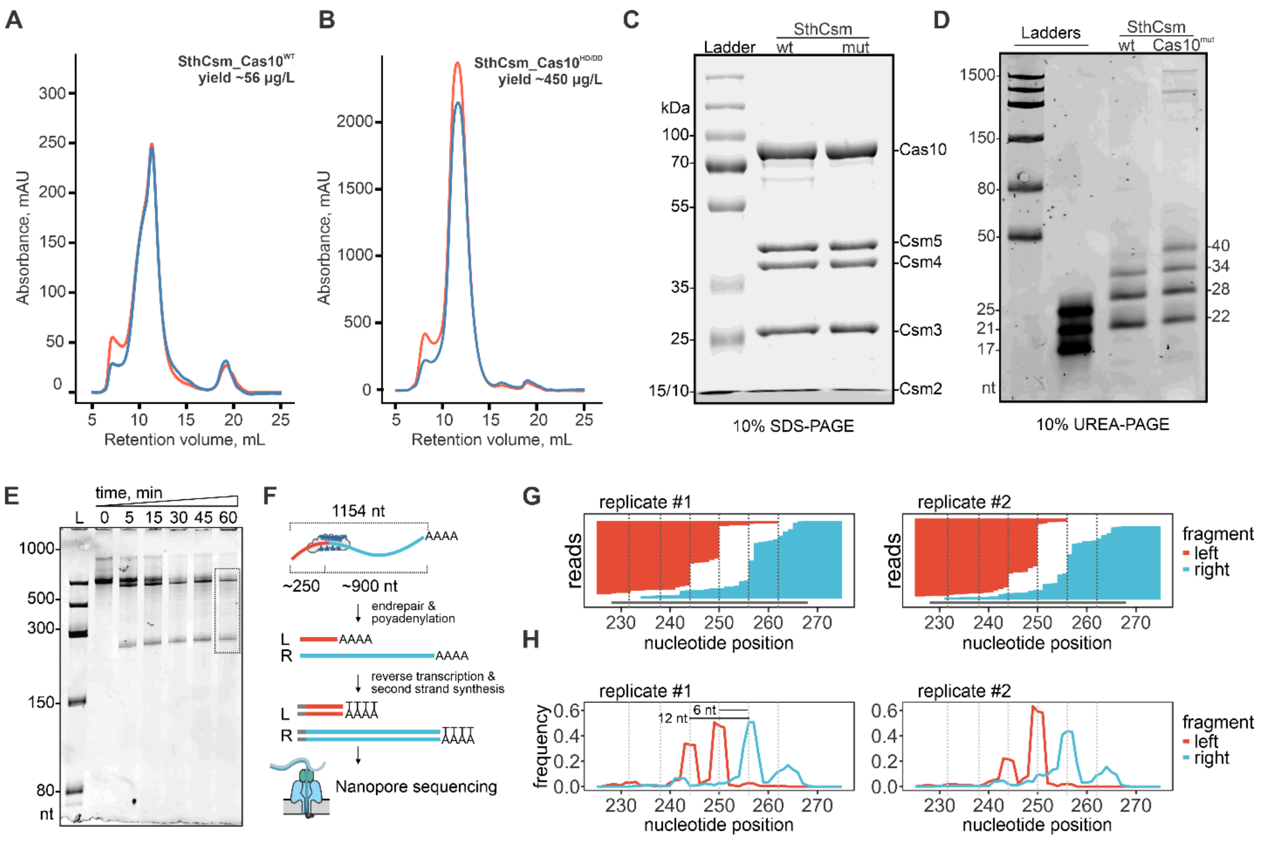


**Fig. S3. Purification and characterization of SthCsm-complexes targeting *gfp* gene in Sindbis virus genome.**

(**A**) Size-exclusion chromatography (SEC) of affinity purified wild-type SthCsm.
(**B**) SthCsm complex with mutated catalytic residues in nuclease (15HD>HA) and polymerase (573GGDD>GGAA) active sites of the Cas10 subunit.
(**C**) SDS-PAGE of purified SthCsm complexes in (A) and (B) (~10 ug/lane).
(**D**) RNAs were extracted from SthCsm complexes and were separated in 12% UREA-PAGE.
(**E**) Cleavage assay with *in vitro* transcribed target RNA. Cleavage reactions were stopped at different time points by adding 2 volumes of phenol:chloroform, extracted, and separated with 8% UREA-PAGE.
(**F**) Diagram of Nanopore sequencing strategy used to determine cleavage sites. L – left cleavage fragment, R – right cleavage fragment
(**G**) Mapping of reads generated by sequencing cleavage fragments produced with SthCsm after 60 min incubation. Red and blue lines show reads generated from left and right cleavage products, respectively. Horizontal black line shows target region complementary to the crRNA. Vertical dotted lines mark predicted Csm3 cut sites.
(**H**) Sequencing reads that end (left fragments, red) or start (right fragments, blue) within the target region were quantified, and resulting frequency was normalized to sequencing depth. Vertical dotted lines mark predicted Csm3 cut sites.

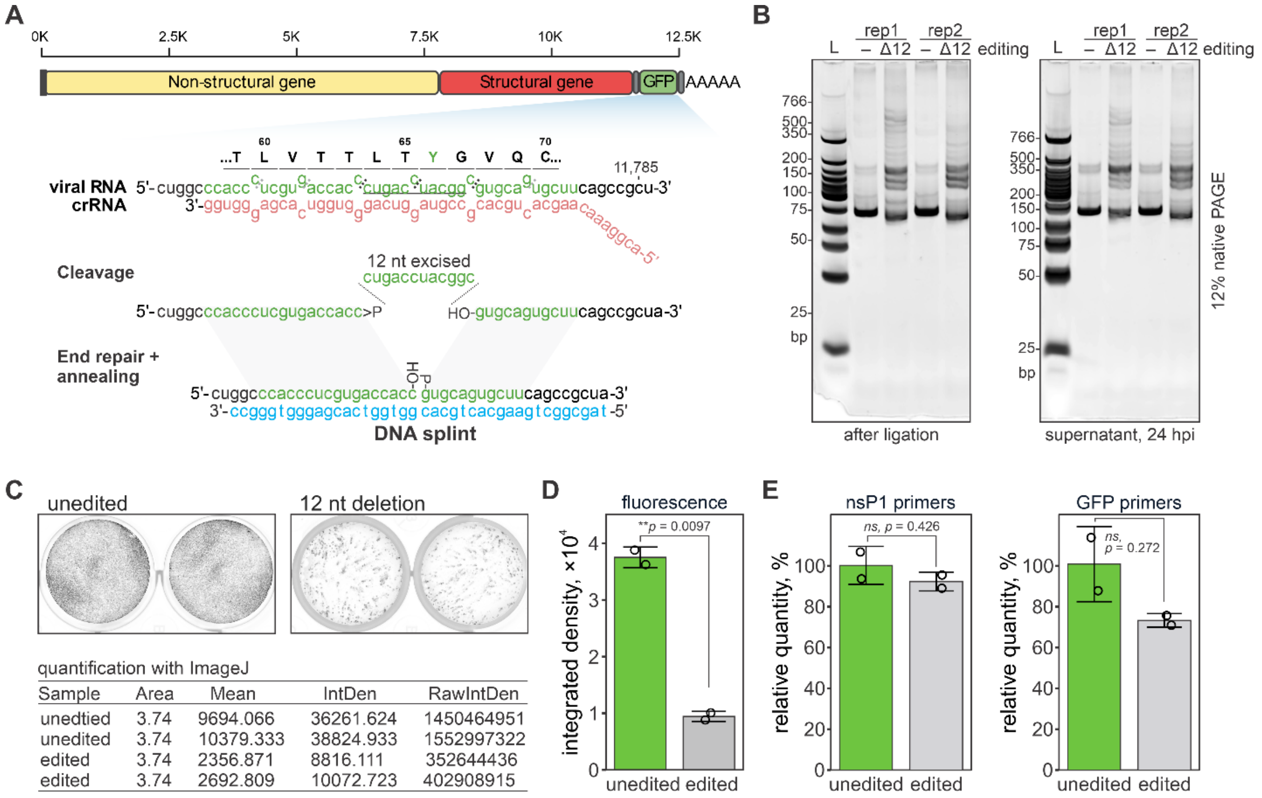


**Fig. S4. CRISPR-based programmed deletions in SINV-GFP.**

(**A**) Experimental design to delete 12 nucleotides in the *gfp* gene of SINV-GFP. The tyrosine in position 66 required for fluorescence is highlighted with green.

(**B**) Related to main Fig. 2. Products generated in RT-qPCR shown in main Fig. 2B and E were analyzed with 12% native PAGE. Two replicates (rep1 and rep2) are shown for the unedited sample (-) and the sample with programmed deletion of 12 nucleotides (Δ12). L – molecular weight ladder.

(**C**) Related to main Fig. 2D. Fluorescence in cells transfected with unedited or edited viral RNA (top) was quantified in ImageJ software after background subtraction. Mean fluorescence, integrated density (IntDen), and raw integrated density (RawIntDen) were calculated.

(**D**) Plot of fluorescence measurements in (C). Bars show the mean of two replicates. Error bars show the mean ± standard deviation. Samples were compared using Welch's unequal variances *t*-test; ** significant difference ($p < 0.01$)

(**E**) Relative levels of RT-qPCR signal in edited vs. unedited sample generated with primers targeting nsP1 (left) or GFP (right) genes in the SINV-GFP genome. Bars show the mean of two replicates. Error bars show the mean ± standard deviation. Samples were compared using Welch's unequal variances *t*-test; ns – no significant difference.

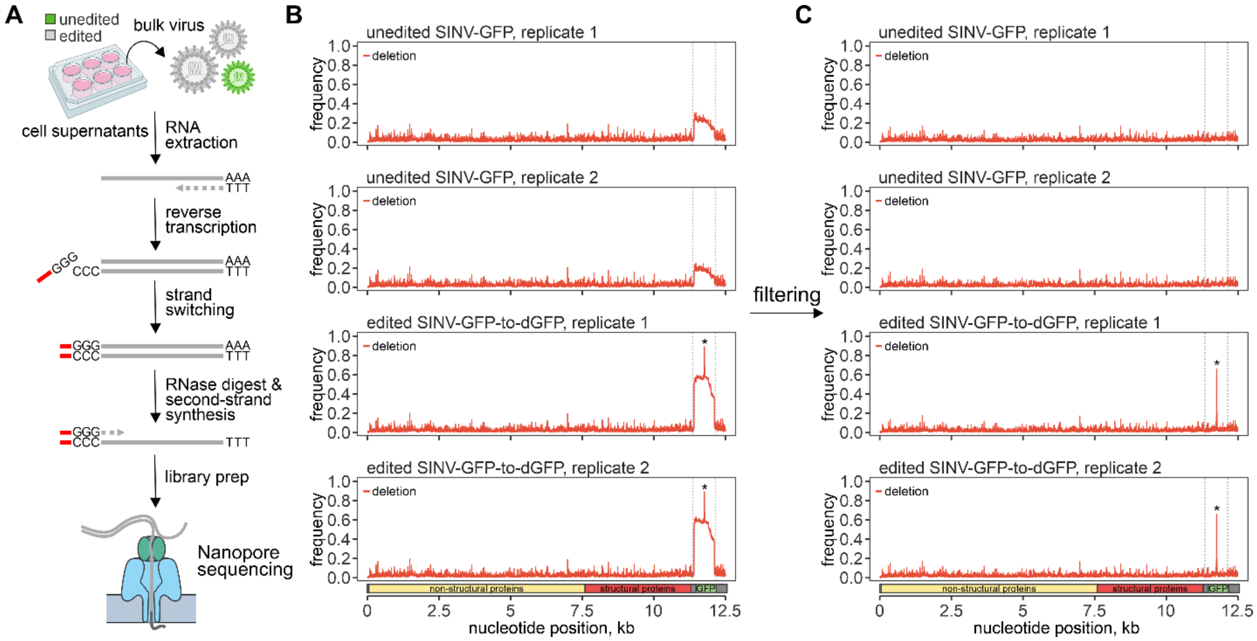


**Fig. S5. Direct cDNA sequencing with Nanopore.**

(**A**) Diagram of the direct cDNA sequencing protocol. RNA from cell supernatants containing bulk virus was extracted and reverse transcribed using poly-T primers. Then, the RNA copy was degraded with an RNase cocktail, and the remaining cDNA first strand was used as a template to synthesize the second DNA strand. The resulting double-stranded DNA was sequenced with Oxford Nanopore.

(**B**) Sequencing reads were aligned to the reference, and deletion frequency was quantified across the entire viral genome. Vertical dotted lines indicate complete *gfp* deletion in the stock virus (unedited). Asterisk (*) marks programmed deletions at the site targeted by the SthCsm complex.

(**C**) Sequencing data in (B) was filtered to remove reads with spontaneous deletions of the entire *gfp* gene.

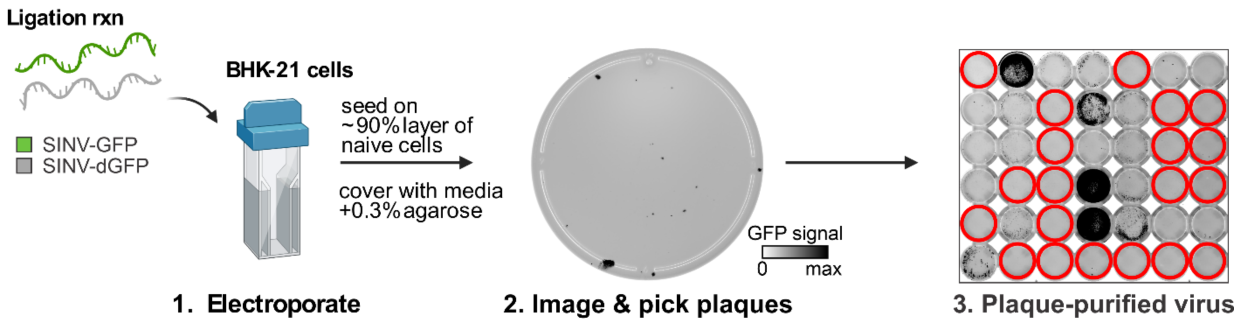


**Fig. S6. Plaque purification of the edited SINV-dGFP virus.**

1) BHK-21 cells were transfected with cleaved, end-repaired, splinted, and ligated RNA. Transfected cells (~10,000 cells) were seeded on a ~90% layer of naïve cells in a 100 mm Petri dish. After 1 h incubation, media was changed for a fresh media supplemented with 0.3% agarose. 2) The next day (16 hpi) Petri dish was imaged with Amersham Typhoon 5 scanner to confirm infection. Then, ~20 hpi viral clones were picked by pushing sterile pipette tip through the agarose overlay. Collected agarose was then used to inoculate naïve BHK-21 cells (~90% confluency) in a 48-well plate. 3) After 24 h incubation, the 48-well plate was imaged with Amersham Typhoon 5 scanner to identify wells without GFP signal but with visible cytopathic effect (red circles).

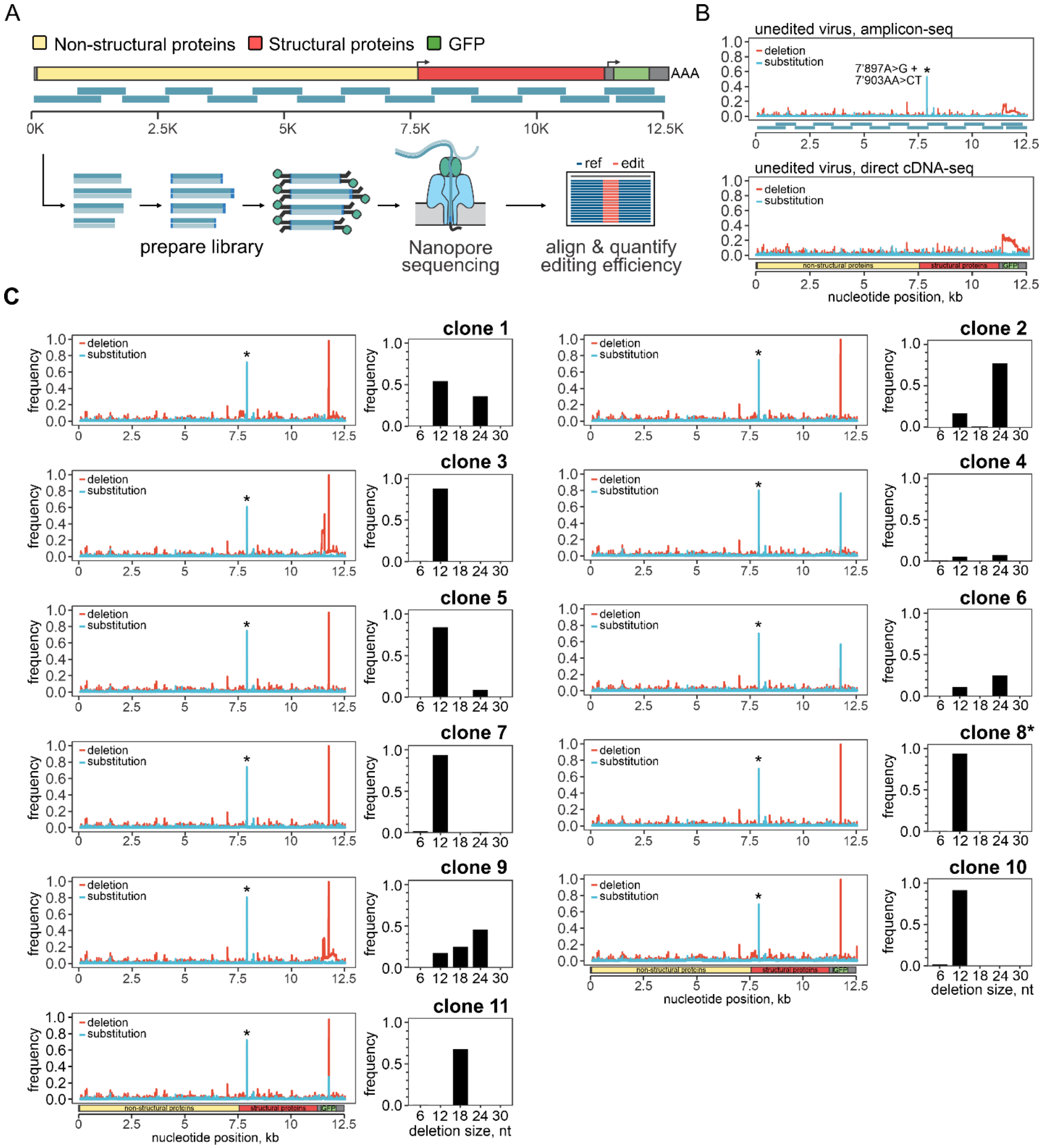


**Fig. S7. Amplicon sequencing of plaque-purified viruses to confirm programmed deletions in the viral genome.**

(**A**) Diagram of the amplicon sequencing strategy used to sequence genomes of plaque-purified SINV. Viral RNA was extracted from supernatants of the infected cell and reverse transcribed. A multiplex PCR was used to amplify ~1 kbp overlapping amplicons that span 97.8% of the viral genome (horizontal teal bars). Resulting amplicons were end-repaired, barcoded, and sequenced with Oxford Nanopore.

(**B**) Unedited bulk virus was sequenced using two sequencing strategies. Sequence variants were identified by comparison to the reference genome. Red shows the frequency of deletions, and blue – nucleotide substitutions. Asterisk (*) indicates 7,897A>G and 7,903AA>CT substitutions that 1) always coincide in the same sequencing reads (~60%), and 2) are not seen in direct cDNA sequencing of the same viral sample (bottom graph). Therefore, we conclude that this substitution is an artifact associated with amplicon sequencing rather than a bona fide genomic mutation.
(**C**) Amplicon-based genome sequencing of 11 plaque-purified viruses (see **Fig. S6**). Sequencing reads were compared to a reference, and all sequence variants were identified and quantified. Red indicates the frequency of deletions, and blue shows nucleotide substitutions. Asterisk (*) marks sequencing artifacts identified in (B) and presumably associated with PCR amplification. Peaks of deletion frequency in the *gfp* gene indicate programmed deletions introduced with CRISPR-based RNA editing (main **Fig. 2** and **Fig. S4**). Bar graphs show the distribution of deletion sizes at the region targeted by SthCsm complex. Sequencing data for clone 8 is shown in the main **Fig 3C**.

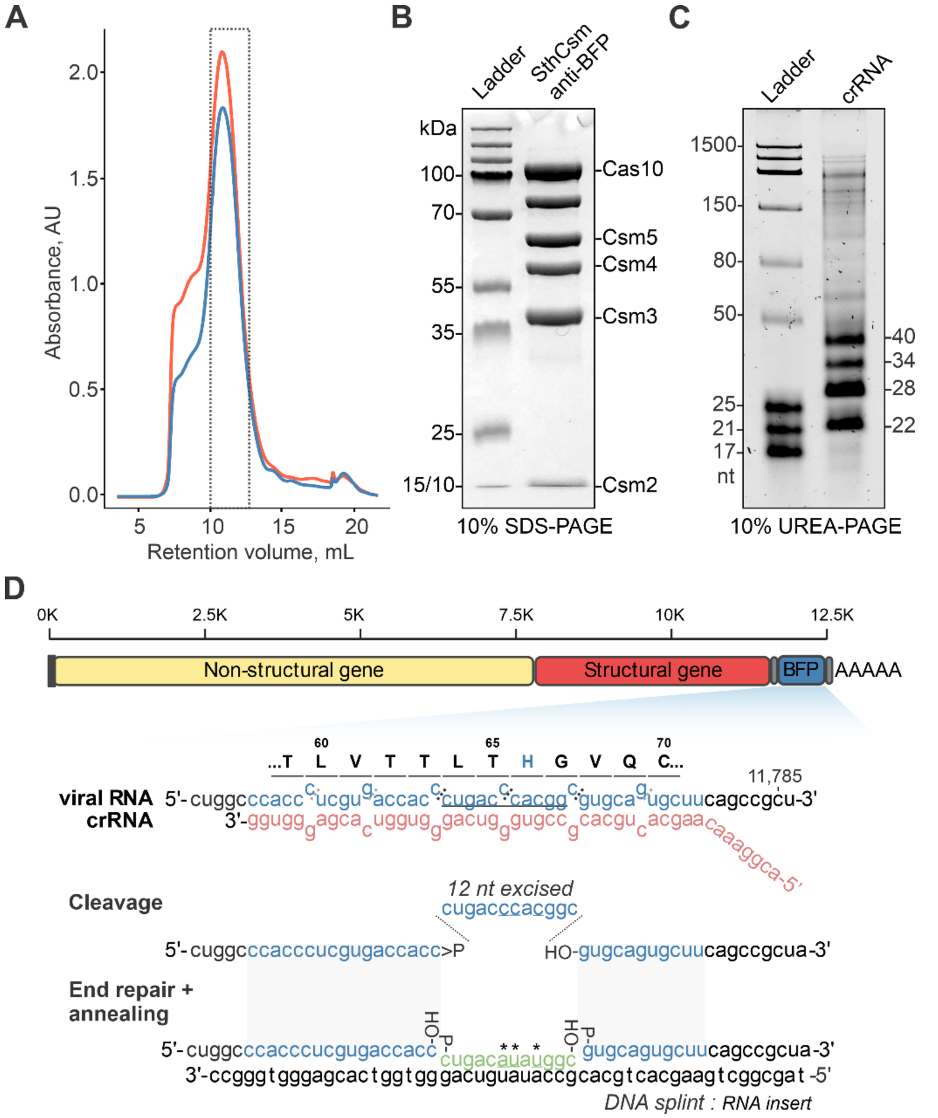


**Fig. S8. Targeting the *bfp* gene in SINV-BFP with SthCsm**

(**A**) Size-exclusion chromatography (SEC) of affinity purified wild-type SthCsm complex targeting *bfp* gene in SINV-BFP. The dotted box shows collected fractions that were pooled together and used for SINV-BFP editing.

(**B**) SDS-PAGE of purified SthCsm complex shown in (A) (~10 ug/lane).

(**C**) RNA was extracted from SthCsm complex purification shown in (A) and separated in 10% Urea-PAGE.

(**D**) Experimental design to substitute 12 nucleotides in the *bfp* gene of SINV-BFP, converting it to GFP. The histidine in position 66 required for fluorescence is highlighted in blue. Asterisk (*) indicates nucleotide substitutions that are introduced into the viral genome through a synthetic RNA insert. The editing replaces the CAC codon with UAU, changing the encoded histidine to tyrosine (H66Y mutation).

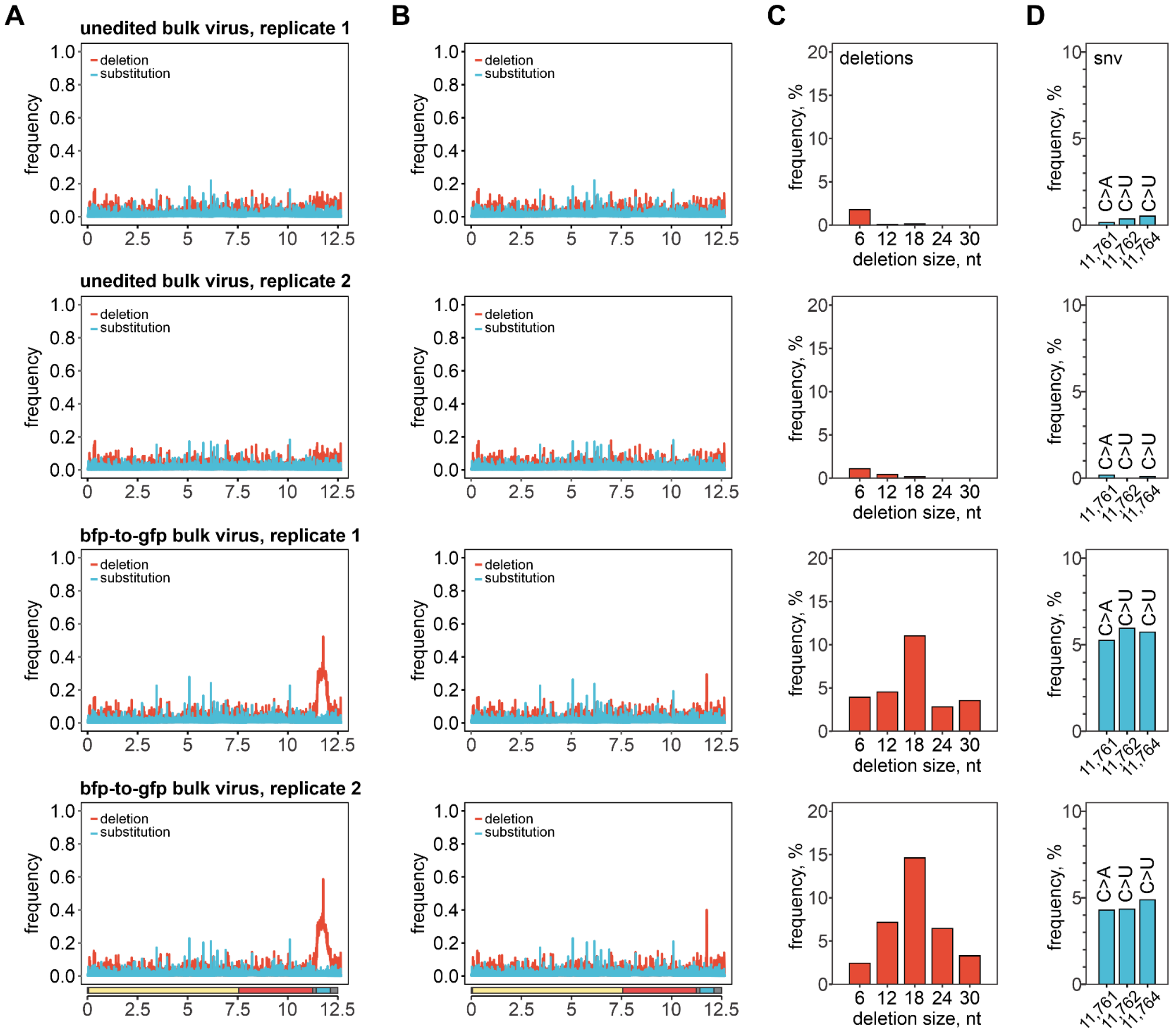


**Fig. S9. Genome sequencing of bulk SINV-BFP-to-GFP viruses**

(**A**) Direct cDNA sequencing of bulk SINV-BFP-to-GFP edited or unedited viruses. Sequence variants in sequencing reads were identified, quantified, and normalized to sequencing depth. Red shows deletions, blue – nucleotide substitutions.

(**B**) Data shown in (A) was filtered to remove reads with complete deletions of the *bfp* gene that are both seen in edited and unedited viruses and accumulate independently of CRISPR targeting during viral passaging.

(**C**) Distribution of deletion sizes at the region targeted by SthCsm.

(**D**) Frequency of programmed substitutions (11,761C>A + 11,762C>U + 11,764C>U) at the target site in the *bfp* gene. Substitution 11,762C>U changes histidine in position 66 of the protein to tyrosine (H66Y mutation), while 11,761C>A and 11,764C>U are silent mutations.

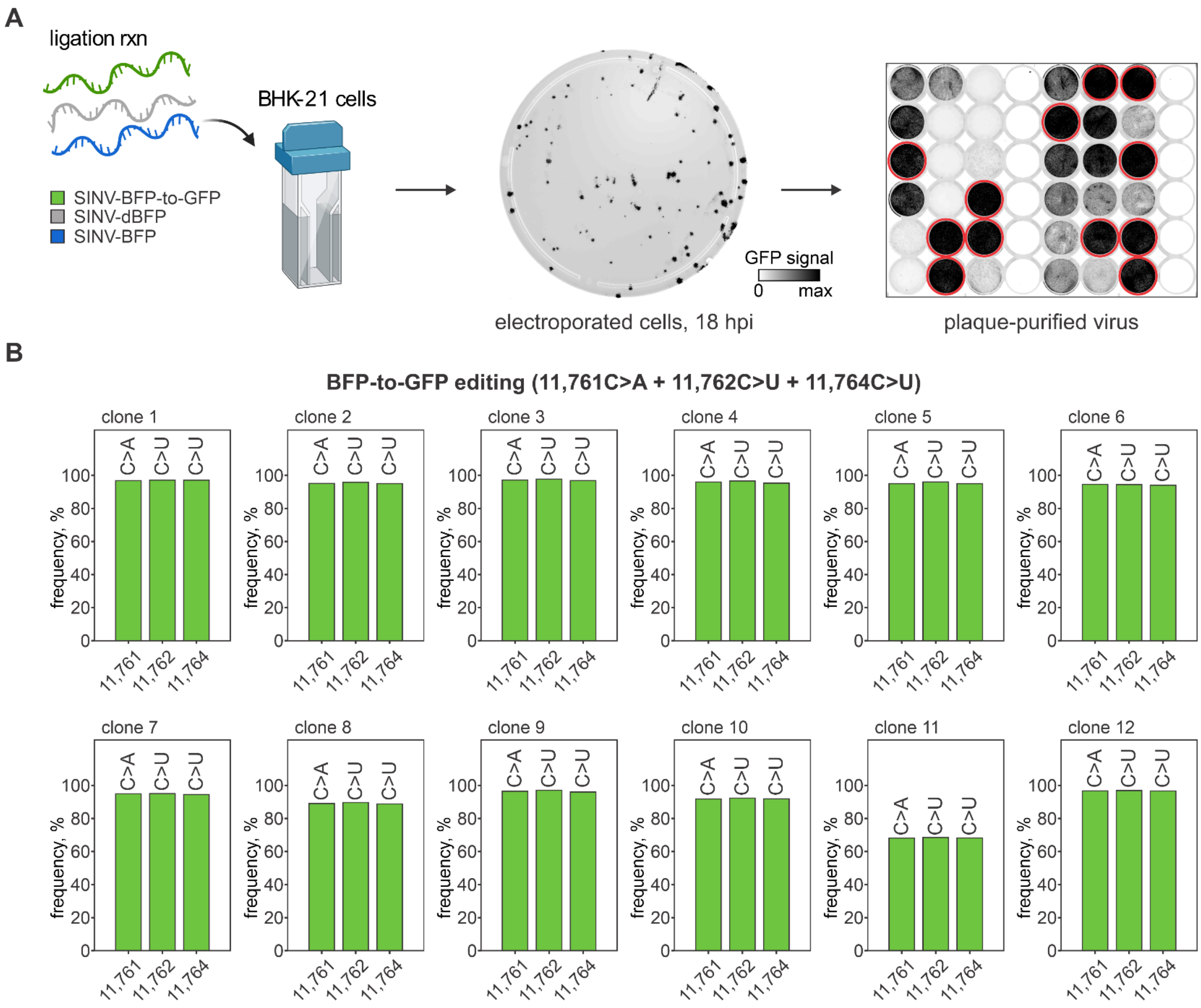


**Fig. S10. Genotyping of plaque-purified SINV-BFP-to-GFP viruses**

(**A**) Plaque purification of SINV-BFP-to-GFP edited virus. Transfected cells (~10,000 cells) were seeded on ~90% confluency layer of BHK-21 cells in a 100 mm Petri dish. After 1 h incubation (37°C, 5% $CO_2$), media was changed for fresh media supplemented with 0.3% agarose. Cells were imaged at 18 hpi. Viral plaques were picked under a standard light microscope and used to inoculate naïve cells in a 48-well plate. After imaging the 48-well plate, 12 wells producing high GFP signal (circled in red) were selected for genome sequencing. (**B**) Frequency of programmed substitutions (11,761C>A + 11,762C>U + 11,764C>U) in the plaque-purified viruses. Substitution 11,762C>U changes histidine in position 66 of the protein to tyrosine (H66Y mutation), converting BFP to GFP, while 11,761C>A and 11,764C>U are silent mutations.

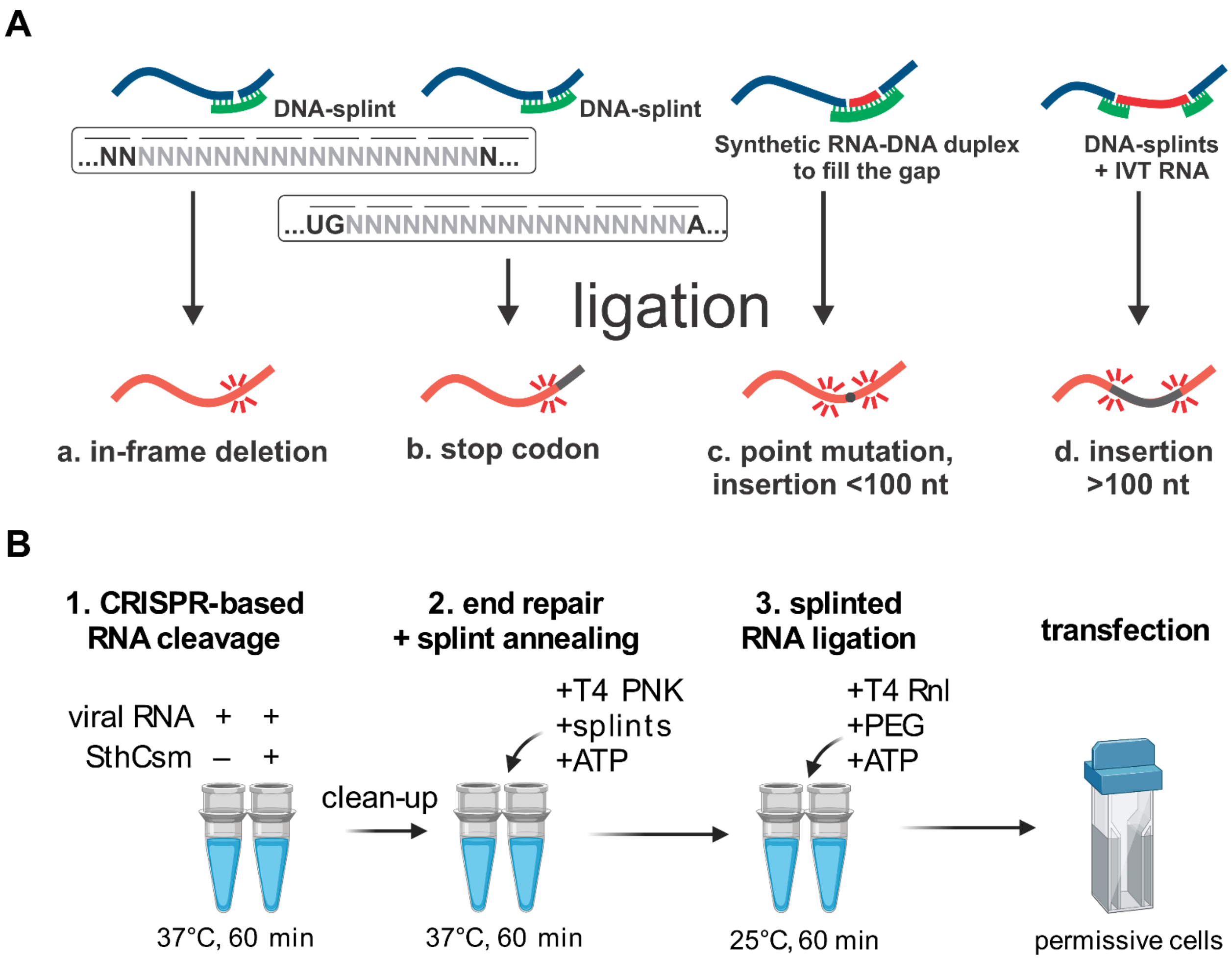


**Fig. S11.**

(**A**) Arsenal of RNA edits enabled by programmable CRISPR-based cleavage and RNA ligation. The 6 nt periodicity of type III CRISPR-based cuts can be leveraged to introduce in-frame deletions (a). "N" represents any nucleotide, the excised section is shown in light gray. Horizontal lines indicate codons. Further, a stop codon can be introduced to knock out the target gene or truncate the protein (b). Dual targeting can delete portions of the genome between cut sites. To insert or substitute a short sequence, synthetic RNA-DNA duplexes with overhangs complementary to the cut site are used (c). In this case, the insertion size is limited by the capacity for the chemical synthesis of RNA (~100-200 nt). For larger inserts, in vitro transcribed (IVT) RNA can be used with splints at 5'- and 3'-ends of the insert (d).

(**B**) Diagram of the CRISPR-based RNA editing protocol developed in this work. The protocol includes only 3 steps that take 3 h of total incubation time and minimal sample handling.

**Table S1.**

Oligonucleotides used for cloning anti-GFP and anti-BGP CRISPR loci

| Name | Sequence 5`-3` |
|---|---|
| Sth_RSR_GFP_F | TAATCCATGGGATATAAACCTAATTACCTCGAGAGGGGACG GAAACAAGCACTGCACGCCGTAGGTCAGGGTGGTCACG |
| Sth_RSR_GFP_R | ATTACAATTGCCCGGGGTTTCCGTCCCCTCTCGAGGTAATTA GGTTTATATCCCACCCTCGTGACCACCCTGACCTACG |
| SDM_BFP_crRNA_F | CAGGGTGGTCACGAGGGTGG |
| SDM_BFP_crRNA_R | ACCCACGGCGTGCAGTGCTTG |

**Table S2.**

DNA splints and RNA inserts used for RNA editing.

| Name | Sequence 5`-3` |
|---|---|
| *Programmed deletions in IVT RNA* | |
| Splint A, 18 nt | TCCGAAGAACGCTGAAGCCAATGTTTGTAATCAGTTCC |
| Splint B, 18 nt | GAAGAACGCTGAAGCGATTTGCGGCCAATGTTT |
| Splint B, 24 nt | GAAGAACGCTGAAGCGGGCCAATGTTTGTAATCAGT |
| *Programmed deletion in SINV-GFP* | |
| Y66_d12_splint | TAGCGGCTGAAGCACTGCACGGTGGTCAC<br>GGGGTGGGCC |
| *Programmed substitution in SINV-BFP* | |
| H66Y_splint | TAGCGGCTGAAGCACTGCACGCCATATGTCAGGGT<br>GGTCAC<br>GAGGGTGGGCC |
| H66Y_insert | /5Phos/CUGACAUAUGGC |

**Table S3.**

Oligonucleotide primers used to generate DNA templates for in vitro transcription of RNA.

| Name | Sequence 5`-3` |
|---|---|
| N0.5_F | TAATACGACTCACTATAGGGCGTGTTGTTTTAGATTTCATC |
| N3-5_R | CACACTGATTAAAGATTGCTATGTG |
| T7_SINV_sgmRNA_F | TAATACGACTCACTATAGTACATTTCATCTGACTAATACTA C |
| SINV_polyA_R | TTTTTTTTTTTTTTTTTTTTTTTTTTTTTTGAAATGTTAAAA ACAAAATTTTGTTG |

**Table S4.**

Oligonucleotide primers used for PCR and qPCR.

| Name | Sequence 5`-3` |
|---|---|
| 11762T>C_F1 | TAATACGACTCACTATAGTACATTTCATCTGACTAATACTAC |
| 11762T>C_R1 | GCACGCCGTGGGTCAGGGTGGTCACG |
| 11762T>C_F2 | GACCACCCTGACCCACGGCGTGCAGTGC |
| 11762T>C_R2 | TTTTTTTTTTTTTTTTTTTTTTTTTTTTTTTGAAATGTTAAAAA CAAAATTTTGTTG |
| CoV2_N_qPCR_F | GGGACCAGGAACTAATCAGAC |
| CoV2_N_qPCR _R | TGTGACTTCCATGCCAATG |
| qSindbisFW4495 (*43*) | TAGACAGAACTGACGCGGACGT |
| qSindbisREV4635 (*43*) | TCCATACTAACTCATCGTCGATCTC |
| qSINV_GFP_F1 | GAGGGCGAGGGCGATG |
| qSINV_GFP_R1 | GCTTCATGTGGTCGGGGTAG |
| nCoV-96_L | GCCAACAACAACAAGGCCAAAC |
| nCoV-96_R | TAGGCTCTGTTGGTGGGAATGT |

**Table S5.**
PCR primers used for amplicon sequencing of SINV-GFP and SINV-BFP genomes.

| Name | Sequence 5`-3` | Pool |
|---|---|---|
| sinv_1K_1v2_L | ACAGCCGACCAATTGCACTA | 1 |
| sinv_1K_1v2_R | AGAGGCTGGGACTTTTACGC | 1 |
| sinv_1K_3_L | CTGAAGAATGCCAAACTCGCAC | 1 |
| sinv_1K_3_R | ATATCCCCTGGCTTCGGCTT | 1 |
| sinv_1K_5_L | CTTGATTTGCAGACGGGGAGAA | 1 |
| sinv_1K_5_R | GCGGTTGTCAAGCAGTTAAGTG | 1 |
| sinv_1K_7_L | GCTTTAGCGGATCGGACAACT | 1 |
| sinv_1K_7_R | ACTGTCCCGTCTACCATATCCAA | 1 |
| sinv_1K_9_L | CGAGTGCGCCTTTGGAGAAATA | 1 |
| sinv_1K_9_R | TTGACGTCGAACAATCTGTCGG | 1 |
| sinv_1K_11v3_L | CTGCCACCATACTGAACCGT | 1 |
| sinv_1K_11v3_R | ATGGTGTACACAGGATGGCG | 1 |
| sinv_1K_13_L | GCGATGAAAGTAGGACTGCGTA | 1 |
| sinv_1K_13_R | GGCTTGGAATGCTCTTTTGCTC | 1 |
| sinv_1K_15_L | CACCTCTAGAATCGCCACCATG | 1 |
| sinv_1K_15_R | GCACCACGCTTCCTCAGAAATA | 1 |
| sinv_1K_2_L | TGTGATACAGTGGTGAGTTGCG | 2 |
| sinv_1K_2_R | GGCATCAGTACTTTAGCGTCGT | 2 |
| sinv_1K_4_L | GCACACAGCCAGTTACAGCTAT | 2 |
| sinv_1K_4_R | ACTGAGTGGTGTTTGAACTGGTT | 2 |
| sinv_1K_6_L | ACAAAACGCCTACCATGCAGT | 2 |
| sinv_1K_6_R | CTTTAGCCTTGGCGGTGGAATA | 2 |
| sinv_1K_8_L | GAACTACTCCGATCCACAGTTCG | 2 |
| sinv_1K_8_R | AGTACTCTGCTGGCGATAACGA | 2 |
| sinv_1K_10_L | CCGAAGAAACCAAAAACGCAGG | 2 |
| sinv_1K_10_R | TCCATGGTGCCTTCTTTAACGG | 2 |
| sinv_1K_12_L | TGGCCTGGAATACATATGGGGA | 2 |
| sinv_1K_12_R | ACGACCTTATGATCGAATGGCG | 2 |
| sinv_1K_14_L | CGCCTCGTCGCTATTAATTATAGGA | 2 |
| sinv_1K_14_R | TACATCGAGTTTTGCTGGTCGG | 2 |
